# Regional and LTP-Dependent Variation of Synaptic Information Storage Capacity in Rat Hippocampus

**DOI:** 10.1101/2022.08.29.505464

**Authors:** Mohammad Samavat, Thomas M. Bartol, Cailey Bromer, Jared B. Bowden, Dusten D. Hubbard, Dakota C. Hanka, Masaaki Kuwajima, John M. Mendenhall, Patrick H. Parker, Wickliffe C. Abraham, Kristen M. Harris, Terrence J. Sejnowski

**Author notes:** For correspondence: (TJS); (KMH); (WCA); (MS).

## Abstract

Connectomics is generating an ever-increasing deluge of data, which challenges us to develop new methods for analyzing and extracting insights from these data. We introduce here a powerful method for analyzing three-dimensional reconstruction from serial section electron microscopy (3DEM) to measure synaptic information storage capacity (SISC) and apply it to data following *in vivo* long-term potentiation (LTP). Connectomic researchers have focused on the pattern of connectivity between neurons. The strengths of synapses have also been studied by quantifying the sizes of synapses. Importantly, synapses from the same axon onto the same dendrite have a common history of coactivation, making them a candidate for measuring the precision of synaptic plasticity based on the similarity of their dimensions. Quantifying precision is fundamental to understanding information storage and retrieval in neural circuits. We quantify this precision with Shannon information theory, which is a more reliable estimate than prior analyses based on signal detection theory because there is no overlap between states, and outliers do not artificially bias the outcome. Spine head volumes are well correlated with other measures of synaptic weight, thus SISC can be determined by identifying the non-overlapping clusters of dendritic spine head volumes to determine the number of distinguishable synaptic weights. SISC analysis of spine head volumes in the stratum radiatum of hippocampal area CA1 revealed 24 distinguishable states (4.1 bits). In contrast, spine head volumes in the middle molecular layer of control dentate gyrus occupied only 5 distinguishable states (2 bits). Thus, synapses in different hippocampal regions had significantly different SISCs. Moreover, these were not fixed properties but increased by 30 min following induction of LTP in the dentate gyrus to occupy 10 distinguishable states (3 bits), and this increase lasted for at least 2 hours. We also observed a broader and nearly uniform distribution of spine head volumes across the increased number of states, suggesting the distribution evolved towards the theoretical upper bound of SISC following LTP. For dentate granule cells these findings show that the spine size range was broadened by the interplay among synaptic plasticity mechanisms. SISC provides a new analytical measure to probe these mechanisms in normal and diseased brains.

## Introduction

In the late 19th century, Santiago Ramón y Cajal proposed that memories are stored at synapses and not through the generation of new neurons (***Ramón y Cajal, 1894***). Since then, there has been an extensive search for synaptic mechanisms responsible for learning and memory. In particular, long-term potentiation (LTP) has become a standard model for investigating cellular, synaptic, and molecular mechanisms of learning and memory. Numerous structural consequences have been shown to accompany LTP. For example, the density and proximity of docked vesicles at the presynaptic active zone area is increased and may explain the enhanced release probability (***Jung et al., 2021***). Dendritic spines and the area of the postsynaptic density (PSD) enlarge at the expense of new spine outgrowth in the mature hippocampus ***(Bourne and Harris, 2011; Bell et al., 2014; Harris, 2020***). Although synaptic plasticity is well-established as an experience-dependent mechanism for modifying these and other synaptic features, the precision of this mechanism is unknown.

The existence of both intrinsic and extrinsic origins of variability and dysfunction of structural modulation (***Kasai et al., 2021***) motivates further exploration of the potential precision with which synaptic strengths can be adjusted. From an information theory perspective, there can be no information stored without precision – the more precise synaptic plasticity is, more distinguishable synaptic states are possible and the greater amount of information that can be stored at the synapses. The synaptic weight is itself the information that is stored at a synapse, and this information is retrieved when synaptic transmission subsequently occurs. Note that synapses are complex dynamical structures and “synaptic weight” is not a single scalar value but rather is a vector composed of intrinsic functional pre- and postsynaptic variables. These variables can be modulated by synaptic plasticity. Included in the vector of variables are the probability of presynaptic vesicular release, number of docked vesicles, number of postsynaptic receptors, and degree of short-term facilitation/depression, to name but a few. Thus over a series of test pulses applied to a given synapse, the resulting train of stochastic excitatory postsynaptic potentials (EPSPs) is expected to vary in amplitude and number of EPSP events from stimulus to stimulus, and vary in a way that is consistent with the synaptic weight vector ***(Kandaswamy et al., 2010; Klyachko et al., 2006)***. Several studies have shown that synaptic strength is highly correlated with dendritic spine head volume ***(Harvey and Svoboda, 2007; Matsuzaki et al., 2004; Harris, 2020, reviewed papers in Yang and Lui, 2022)***. Pairs of dendritic spines on the same dendrite that receive input from the same axon (SDSA pairs) occur naturally in the brain and are expected to have experienced the same activation histories ***(Harris and Sorra, 1998; Kumar et al., 2020)***. Hence, synaptic weight precision can be estimated by measuring the difference between the spine head volumes of SDSA pairs, and this information can then be used to calculate the number of distinguishable synaptic states in a synaptic pathway.

In previous studies ***(Bartol et al., 2015, Bromer et al., 2018)***, signal detection theory was used to estimate the number of distinguishable synaptic strengths with an assumed signal-to-noise ratio of 1 across the range of spine head volumes. Signal detection theory determines the probability of detecting a signal in noise depending on the overlap between the two distributions. A threshold for the amount of overlap determines the separation between them. In its application to identify distinguishable synaptic strengths, we chose an overlap threshold of 31%, so that an ideal observer would correctly assign a given sample to the correct distribution 69% of the time. Outcomes from these analyses of CA1 pyramidal cell apical dendrites revealed a remarkably high precision with 26 Gaussian distributions spanning the range of SDSA pairs (***Bartol et al., 2015***). However, signal detection theory has limitations. For example, if a single large spine head volume (e.g., a 0.55 *µm*^3^ spine head found in another dataset, ***Harris and Stevens, 1989***) were added to the dataset, we would predict 39 distinguishable Gaussians spanning the range, an increase of 50% in the number of states. In ***Bartol et al. (2015)*** there are interval gaps with Gaussian distributions across a range of spine head volumes without spine head volumes in the reconstructed tissue. Furthermore, this previous method assumed an arbitrary threshold for the amount of overlap allowed between the consecutive Gaussians. Finally, the method did not take into account the actual frequency of spine head volumes in the distinguishable states.

We used information theory to calculate the number of bits of Shannon information stored in synaptic weights to quantify empirically the synaptic information storage capacity (SISC). Information theory is based on the distinguishability of messages sent from a transmitter to a receiver. In the case of synaptic weight, the postsynaptic dendrite or soma is the receiver, which has to distinguish the strengths of messages coming from discrete synaptic inputs. The distinguishability of the received messages depends on the precision with which synaptic plasticity sets the strength of individual synapses. Statistics and information theory allowed us to quantify the precision and information capacity of the message and how it changes following LTP induction. The analysis of precision in our new method starts with measuring the coefficient of variation (CV) of SDSA pairs, the same starting point as in ***Bartol et al. (2015)***. The new method, however, performs non-overlapping cluster analysis to obtain the number of distinguishable categories of spine head volumes in the reconstructed volume using the precision level estimated from the CV of the SDSA pairs. In the new SISC analysis, the number of distinct synaptic states defined by the individual clusters converges toward the true number of states as the number of spine head volumes increases and the true shape of the distribution is sampled. There are more distinguishable states with greater Shannon information when the precision is higher.

Comparison of the new SISC measurements with the previous results demonstrates that the new method is more robust to outliers and, importantly, can reveal gaps and variation in the shape of the distribution. In contrast, with signal detection theory the gaps were filled in with Gaussians in the absence of any data. SISC was applied to synapses in control hippocampal area CA1 that was perfusion-fixed *in vivo*, and in the dentate gyrus perfusion-fixed at 30 minutes and 2 hours after the induction of unilateral LTP compared to the contralateral control hemisphere *in vivo*. The results revealed robust differences between the brain regions and across conditions following synaptic plasticity.

## Results

### Induction of LTP in the Dentate Gyrus

We analyzed datasets of 3D reconstruction from serial section electron microscopy (3DEM) containing perforant path synapses in the middle molecular layer (MML) of the dentate gyrus for inputs arising from the medial entorhinal cortex. Data were collected from the stimulated hippocampus of two rats at 30 minutes and two rats at 2 hours post-induction of LTP, with the hippocampus in the opposite hemisphere serving as the control. All experiments were conducted in the middle of the animals’ waking (dark) period to control for variation due to the circadian cycle (***Bowden et al., 2012***). Analysis of the 30 min control and LTP datasets using our previous signal detection method (***Bartol et al., 2015***) was published in ***Bromer et al. (2018)***. Although the stimulus protocol is known to induce multiple forms of plasticity including LTP, heterosynaptic LTD ***(Abraham & Goddard 1983, Bowden et al 2012)***, and EPSP-spike potentiation (***Abraham et al, J Physiology, 1985)***, for simplicity we refer to the stimulated groups as 30 min LTP and 2 hr LTP, as this is what was measured electrophysiologically on the stimulated pathways.

We used previously described methods to induce LTP in the MML of freely moving rats (***Bowden et al., 2012***). Briefly, stimulating electrodes were surgically implanted in both the medial and lateral perforant paths of the LTP hemisphere, and an additional stimulating electrode was implanted in the medial path of the control hemisphere. Field potential recordings were made using electrodes placed bilaterally in the dentate hilus. Animals were allowed to recover for two weeks prior to producing LTP or control stimulation during the animals’ dark (waking) part of the circadian cycle. LTP was induced by 50 trains of unilateral delta-burst stimulation (DBS) to the medial path electrode and then recorded for either 30 min or 2 hr, timed from the beginning of the delta-burst stimulation. Relative to the two control hemispheres, the LTP hemispheres showed an average of 41% potentiation in the MML for the 30 min experiment (*Fig. 1A, B*). In the 2 hr experiment, there was an average of 37% LTP for the two animals (***Fig. 1***C).

**Fig. 1:**
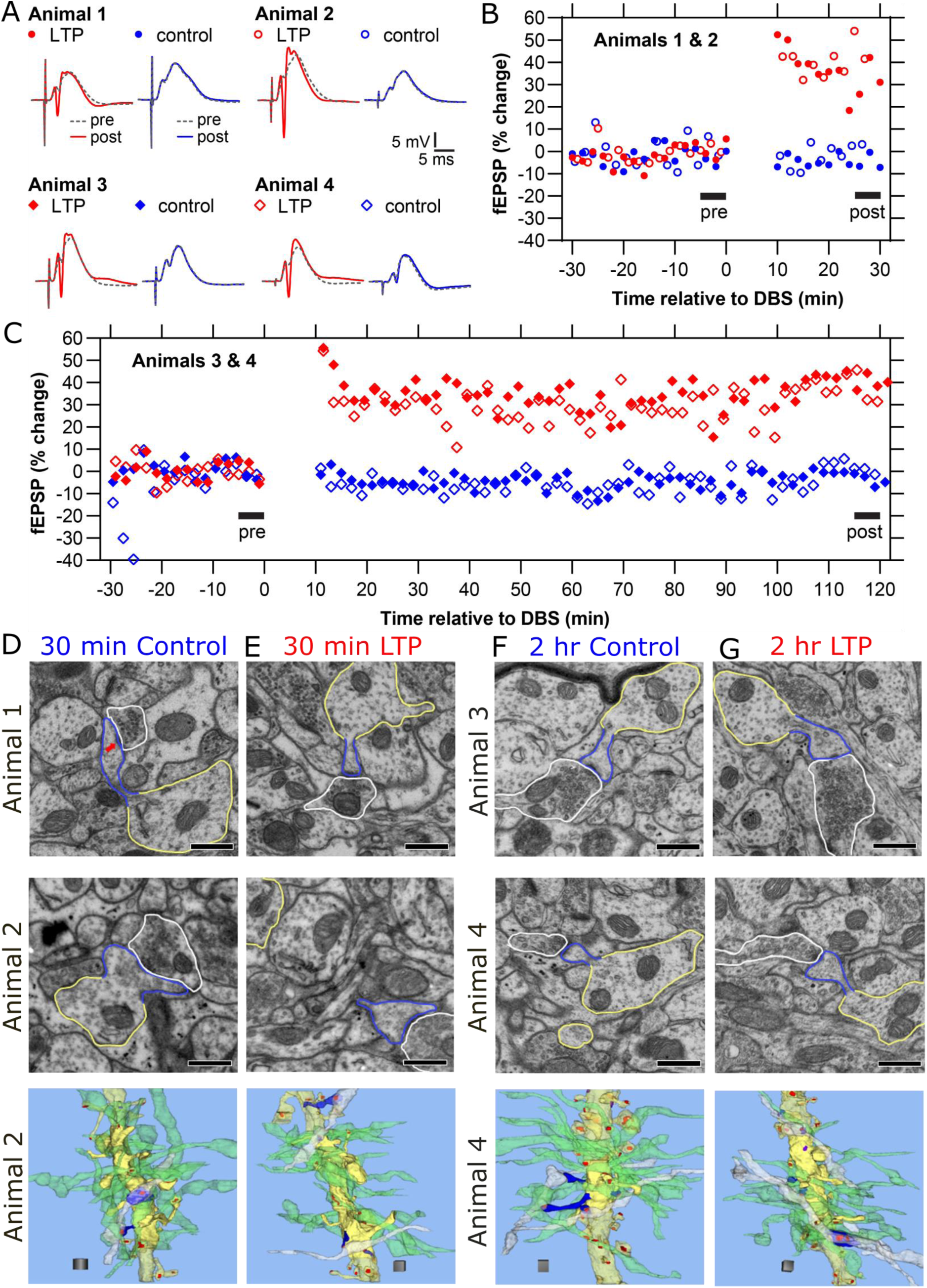
LTP and control responses monitored for 30 min and 2 hours prior to preparation for 3DEM, and representative dendrites from the control and LTP hemispheres in MML. (A) Representative waveforms from baseline responses (dotted, pre) superimposed on responses following delta-burst stimulation (solid, post) in the LTP (red) or control (blue) hemispheres. (Unique symbols are indicated for each animal and plotted in B and C). (B) Change in fEPSP slopes relative to baseline stimulation in the LTP (red) or control (blue) hemispheres monitored for 30 minutes prior to fixation. The average change relative to baseline stimulation in fEPSP response was 34% and 48% at 30 minutes post-LTP induction and 0% for controls. (C) Change in fEPSP slopes relative to baseline stimulation in the LTP (red) or control (blue) hemispheres monitored for 2 hours prior to fixation. The average change in fEPSP slopes relative to baseline stimulation was 41% and 34% for the LTP (red) and 0% for control (blue) hemispheres. (D-G) Example electron micrographs (with red arrow in **Fig. 1**D (first row) pointing to the PSD) and 3D reconstructions in the control and LTP hemispheres as indicated for each of the 4 animals. (Scale bars = 0.5 *µm*.) Bottom row illustrates representative dendrites from control and LTP conditions in Animals 2 and 4 with segment lengths across the row of 9.25, 10.62, 9.44, and 11.33 *µm*, respectively. Axons synapsing on 15 spines along the middle of the dendrite (solid yellow) were analyzed for presynaptic connectivity. Most of the axons (green) made synapses with just one dendritic spine, and some axons (white) made synapses with two dendritic spines (blue). Thus, the white axons illustrate the SDSA pairs. The dendritic shaft and spines occurring along the rest of the reconstructed dendrite are illustrated in translucent yellow. All excitatory synapses are illustrated in red, and the inhibitory synapses in purple. Scale cube = 1 *µm*^3^. ***Supplementary Videos 1-4*** *for 3D illustration of* ***Figs. 1****D-G* are provided.

Serial electron micrographs and 3D reconstructions were prepared from the control (***Fig. 1***D) and 30 min LTP ***(Fig. 1***E) hemispheres of two animals, and the control ***(Fig. 1***F) and 2 hr LTP ***(Fig. 1***G) hemispheres of the other two animals. Three-dimensional reconstructions were made for all of the dendritic spines and synapses occurring along three dendritic segments from each of the control and LTP hemispheres for a total of 24 dendrites and 862 dendritic spines. Axons that were presynaptic to at least 1 of 15 dendritic spines located along the middle of the dendritic segment were traced to determine whether they made more than one synapse along the same dendrite, and thus formed SDSA pairs. All 3D reconstructions and measurements were obtained blind as to condition or animal.

### Comparison of Spine Sizes of 30 min and 2 hr LTP Conditions to Control Stimulation

We analyzed the 4 dentate gyrus MML datasets to see how LTP at 30 min and 2 hours post-induction affected spine head volumes. ***Fig. 2*** compares the spine head volume histograms before and after the induction of LTP. Control histograms from the unstimulated hemisphere (***Fig. 2A***, 2B) are presented above their corresponding LTP histograms ***(Fig. 2***C, ***2***D). The differences between the LTP and the control histograms revealed increases in the numbers of both large and small synapses at both time points (***Fig. 2***E). These findings suggest potentiation of stimulated synapses and concurrent depression of presumptive non-stimulated synapses. However, by 2 hours the peaks and troughs shifted such that the increase in smaller spines was significantly reduced and the increase in larger spines was significantly consolidated (***Fig. 2***F). We designed a statistical test (see methods for details) to test if the observed summed differences between the frequency of 2 hr LTP and that of 30 min LTP (across equal width bins on a log scale for spine head volumes) is significant. We found the significance of the observed differences in ***Fig. 2***F. The P value = 6 ×10−4 for the observed bump in ***Fig. 2F*** for spine head volumes greater than median value (vertical dashed line), and P value = 0.0015 for the observed troughs smaller than median. The same test was performed on all observed peaks and troughs of the trajectories in ***Fig. 2***E,F and are significant (P Value < 0.005).

**Fig. 2:**
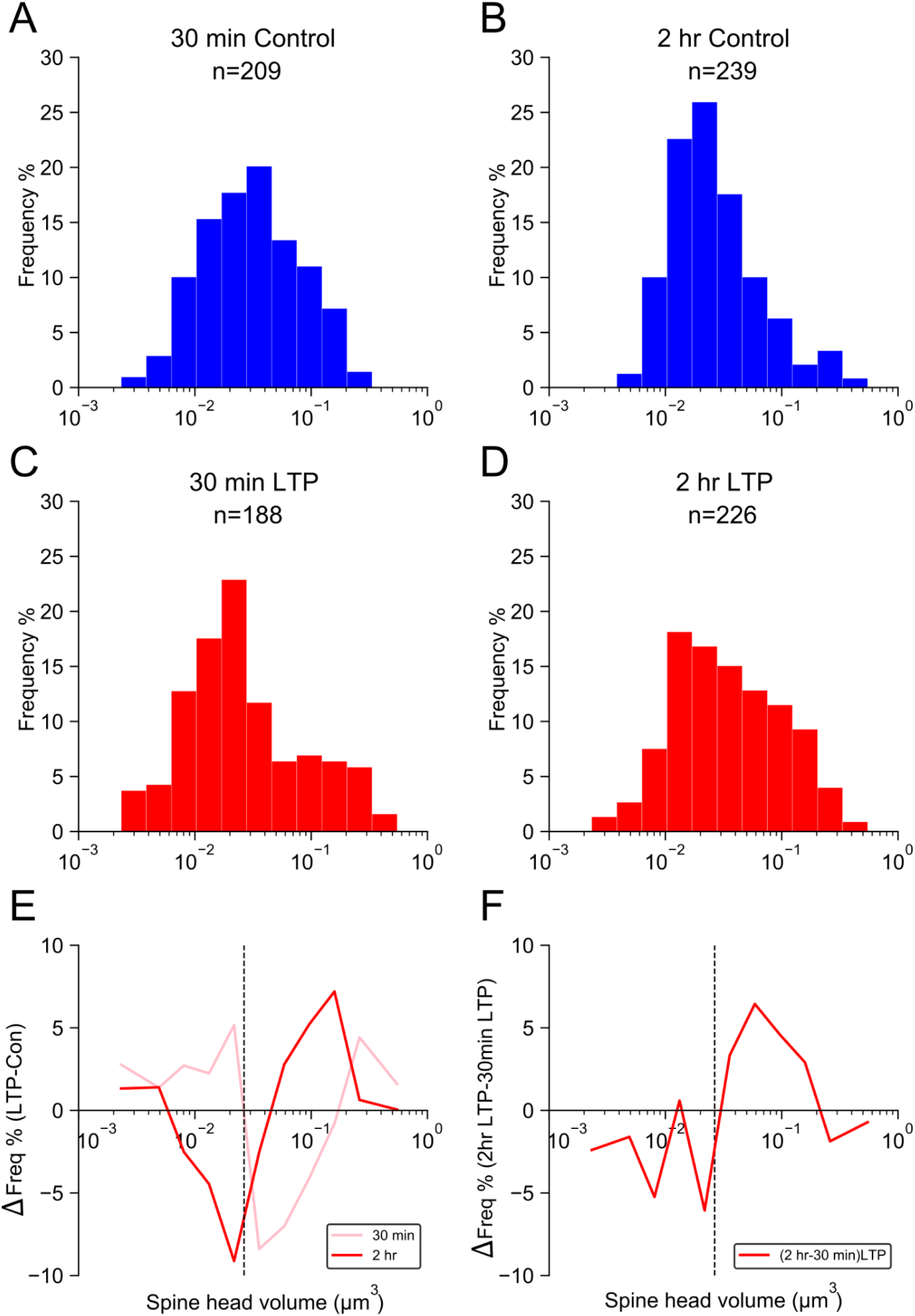
Change relative to control hemispheres in the distribution of spine head volumes at 30 min and 2 hr after the induction of LTP. (A-D) Frequency distributions of spine head volumes (on log scale) from control and LTP hemispheres as indicated. (E) Difference between the frequency of spine head volumes in control and LTP conditions (i.e., LTP-control) at 30 min. (F) Difference between the frequency of spine head volumes in 30 min LTP (C) and 2 hr LTP (D) conditions. The differences in the summations of the representative frequencies (the test statistic) in the observed peaks and troughs in both Fig. 2E*, F* were tested for significance by hypothesis testing with nonparametric bootstrap (see methods). The vertical dash line is the median of combined dentate gyrus spine head volumes. All observed peaks and troughs in both Fig. 2E*, F* were found to be statistically significant (P < 0.005 for all peaks and troughs).

This finding means that by 2 hours the largest synaptic spines were markedly increased in frequency. To evaluate whether the increase in both small and large spine heads resulted in a balanced total synaptic input, we performed an unbiased dendritic segment analysis based on the reconstructions of the intermediate portions of the dendritic segments for all 24 reconstructed dendrites (***Fig. 1****D, E*, *F*, G, Animal 2, solid yellow). None of the findings could be explained by changes in the number of spines or axons per micrometer length of dendrite that were similar between the control and 30 min and 2 hr LTP dendrites (***Fig. 3****A, C*). Furthermore, the summed asymmetric synaptic area across all synapses per micrometer of dendritic length was constant across the control and 30 min and 2 hr LTP conditions (***Fig. 3****B, D*). Thus, the enlargement of some spines was counterbalanced by shrinking of others, and the summed synaptic input remained constant along these local stretches of dendrite across time and conditions, consistent with previously published results in CA1 ***Bourne and Harris (2011)***.

**Fig. 3:**
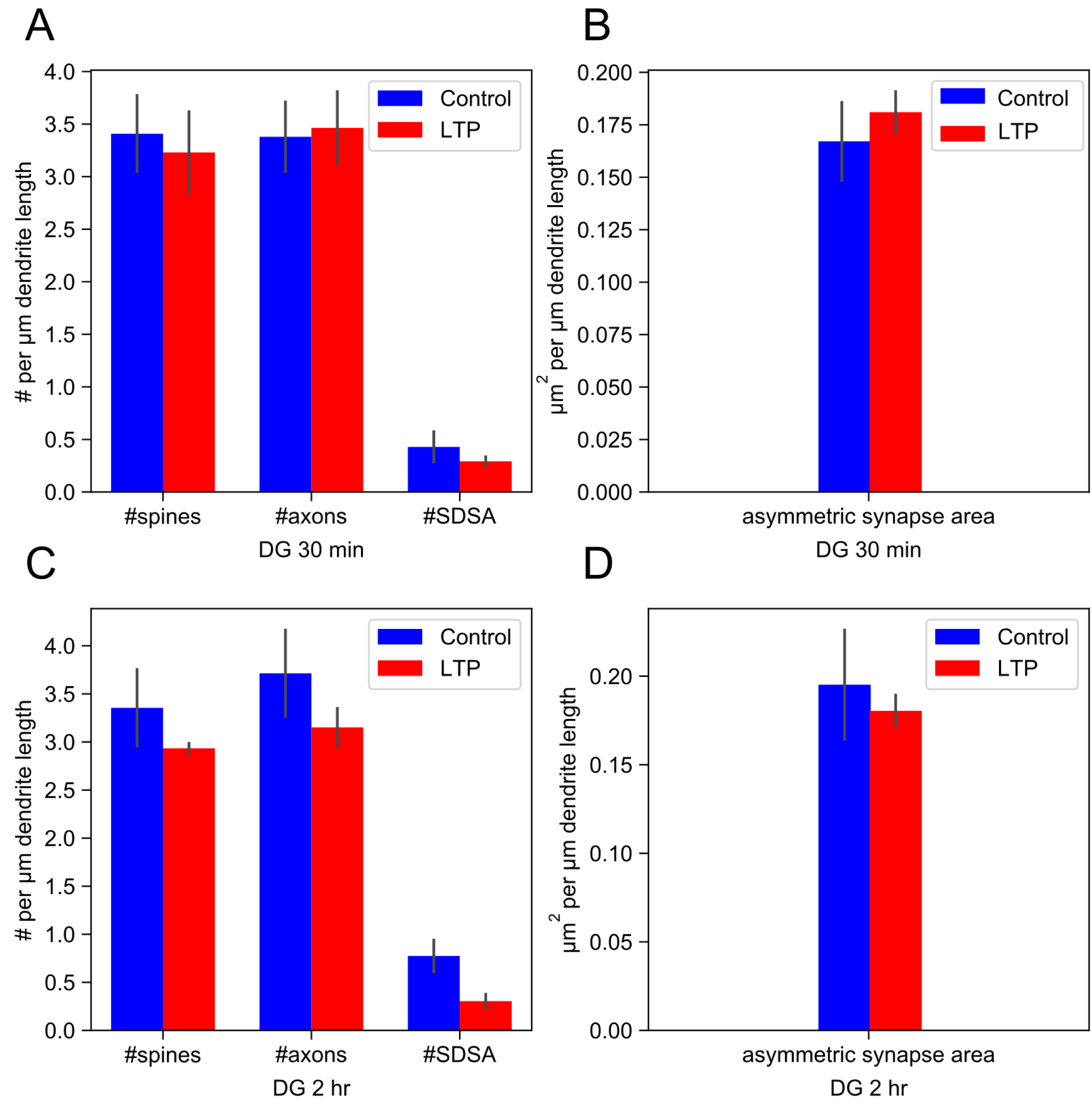
LTP does not change the number of spines, axons, or the summed synaptic area per unbiased length of dendrite in the dentate gyrus. (A) (C) Bar plot of the number (per micrometer length of dendrite, mean ± SEM) of spines, axons, and axons participating in SDSA pairs. There is no significant difference between control (ctrl) and LTP hemispheres for 30 min and 2 hr except for the SDSA counts. (B) (D) The total asymmetric synapse area (based on the PSD area per dendrite micrometer), including spines and asymmetric shaft synapses) was not significantly different between the two LTP conditions relative to control.

We repeated this analysis for each rat (**Supplementary Fig 1**). The peaks and troughs of differences between the LTP and the control histograms at both time points were also significant, confirming the robustness of these findings.

### Precision Analysis

Precision is defined as the degree of reproducibility of a measurement and is often mistaken for accuracy, which is defined as the deviation of the average measurement from a reference value (**Supplementary Fig. 2**). The CV shown in equation (eq) 1 is a statistic that measures the variations within a sample, defined by the standard deviation (σ) eq 2, normalized by the mean of the sample (μ), making it a useful metric for measuring precision. Here we used *N* = 2 in eq 2 because we analyzed SDSA pairs.

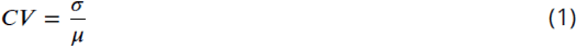

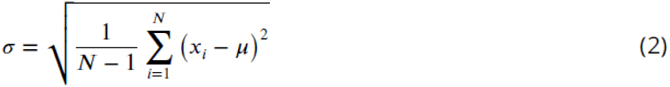

Precision is a key factor for discovering the number of distinguishable states for spine head volumes. The precision is used by the clustering algorithm (see Methods “Clustering Algorithm”) to assign each spine head to its appropriate bin. To estimate the precision, first we determined that the measurement error among 4 investigators who independently measured all of the spine head volumes in the CA1 dataset (**Supplementary Fig. 3**) was smaller than the intrinsic variability of the measured spine head volumes of the SDSA spine pairs. Then we could measure the precision of spine head volumes within the SDSA pairs to estimate the precision of synaptic plasticity. To do this we calculated the CV of all SDSA pairs in each of the 5 datasets ***(Fig. 4)***. None of the correlations between the CV values and mean spine head volumes for the SDSA pairs within each condition were significant. The CVs among SDSA pairs vary from pair to pair by over an order of magnitude within datasets for control and 30 min and 2 hr post LTP induction conditions (Fig. 4), but with no significant trend from the smallest to the largest spine head volumes. These outcomes suggest that the synaptic plasticity based on co-activation history among small spines is as precise as it is for large spines for both control and LTP conditions. It is important to note that the estimate of precision of synaptic plasticity provided by the SDSA pairs applies to all synapses of the same type in a given brain region, and not just SDSA pairs.

**Fig. 4:**
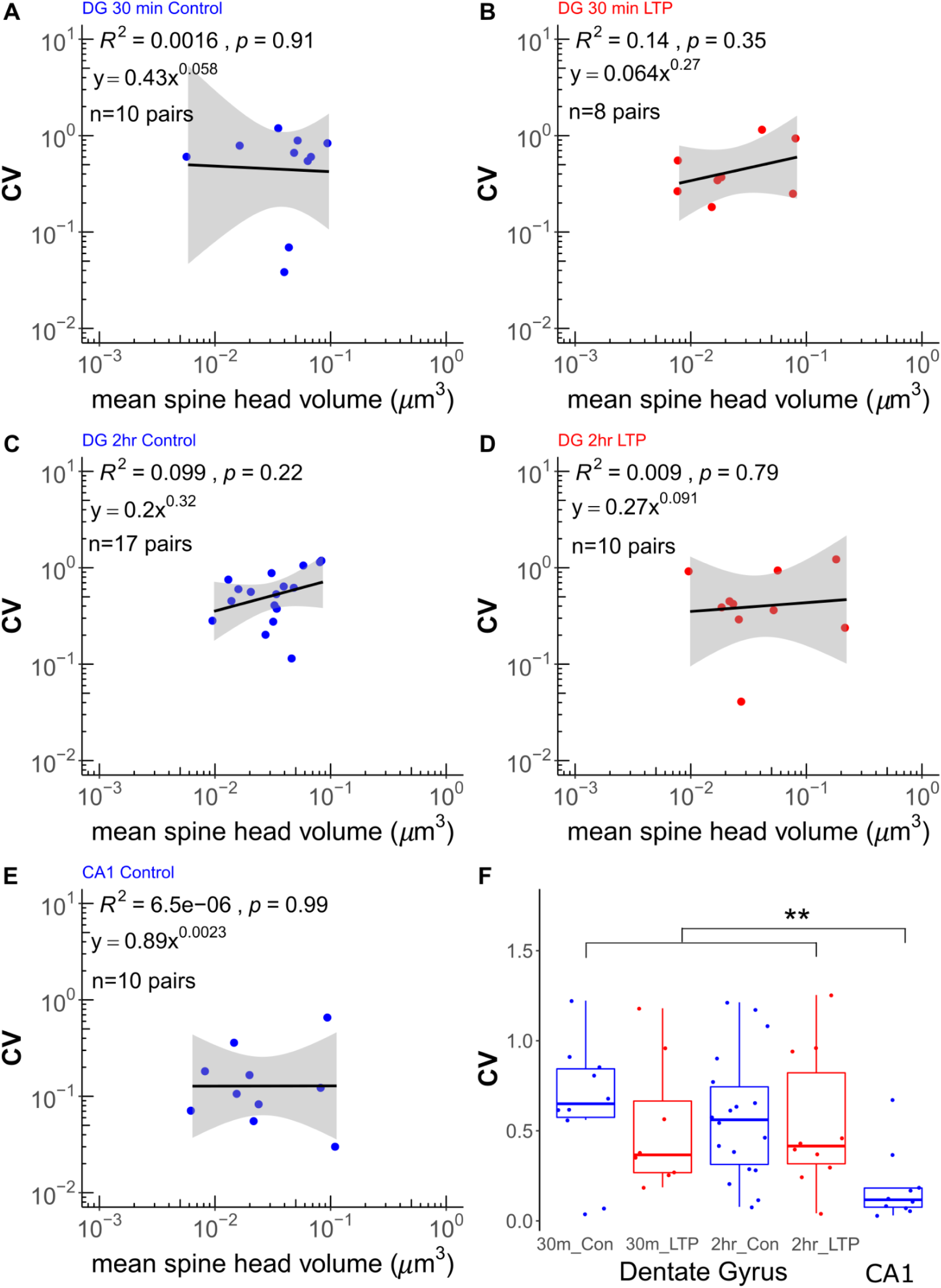
Analysis of synaptic precision based on CV of SDSA pairs across brain regions and plasticity. (A-E) Same-dendrite same-axon (SDSA) pairs were analyzed from each dataset. The regression line, *p* value, and *R*^2^ for the CV of n SDSA pairs are shown for each indicated condition and hippocampal region (The lowest CV of Fig. 4 *C, n=18* is treated as an outlier and removed). The gray region is the 95% confidence interval for each regression line. The Y axis is the CV for each SDSA pair depicted by blue and red dots in control and LTP conditions, respectively. The X axis shows the mean value of the spine head volumes, on a log scale, for each SDSA pair. (F) Summary medians and interquartile ranges (See methods for details) for the dentate gyrus and CA1 datasets combined (overall Kruskal-Wallis *p*-value=0.00085) show there is a statistically significant difference between the CV of the CA1 SDSA pairs and those in dentate gyrus (asterisks represent significance of the p (*<0.05; **<0.01)). To test differences between the control and LTP CVs within the dentate gyrus, the 2 lowest CV values from dentate gyrus 30 min control and 2 hr control are treated as outliers and removed. The one factor KW test (on the first four columns) showed no significant difference between the four dentate gyrus conditions (*p*=0.16). Interestingly, the p value of the one factor KW test between the CV of the dentate gyrus control (30 min and 2 hr combined) and LTP (30 min and 2 hr combined) is not significant (*p*=0.06), but is a trend that could become significant with more data points. Thus, the precision level was significantly higher in area CA1 than dentate gyrus, but not significantly different across the dentate gyrus control and LTP conditions.

In addition, the difference between the CV of SDSA pairs in CA1 was much less (***Fig. 4***E) than in the combined dentate gyrus datasets, which did not differ from one another (***Fig. 4***F). Thus, the CV of the SDSA pairs did not differ significantly across the dentate gyrus MML conditions, but did differ significantly between the two hippocampal regions. The median CV value establishes the precision level of the sets of SDSA pairs in each of the 5 datasets and is used below for cluster analysis and calculation of the number of distinguishable synaptic strength levels. The rationale behind using median CV as a constant threshold for clustering spine head volumes across the range of spine head volumes is our observation that small spines are as precise as large spines for both control and LTP conditions.

### Measurement of the Distinguishable State Distribution

To introduce and compare the performance of our new method for measuring the distinguishable state distribution, we reanalyzed the CA1 dataset that was previously analyzed with signal detection theory (***Bartol et al., 2015***). A total of 288 spine head volumes were fully contained within a 6 × 6 × 5 *µm*^3^ CA1 neuropil volume (***Fig. 5***A). Signal detection theory revealed 26 distinguishable Gaussian distributions with equal CV (inset, ***Fig. 5***B) when assuming an overlap of 31% (***Fig. 5***B). This amount of overlap is equivalent to assuming a signal-to-noise ratio = 1 and a 69% discrimination threshold common in psychophysics (***Schultz, 2007***). Our new clustering method, based upon the median CV of 0.12 ± 0.046 of the SDSA pairs without any assumptions regarding the signal-to-noise ratio (Algorithms 1 and 2, methods), placed the CA1 spine head volumes into 24 distinguishable categories (***Fig. 5***C). The histogram of spine head volumes on a log scale is shown in the inset on panel 5C. The width of the bins was determined by the protocol used for the precision analysis ***(Fig. 4)***, and is equal to the median CV of the SDSA pairs. The resulting distribution represents the frequency of spine head volumes within their respective categories of distinguishability. Visualizing the data in this way displays the number of spines observed in each of the distinguishable clusters of spine head volumes and the vacant spaces across the distribution of sizes. Note that the shape of the distribution of distinguishable states is not expected to be the same as the histogram of spine head volume frequency. Also, note that bins having a width of the same CV appear as equal width bins in log scale.

**Fig. 5:**
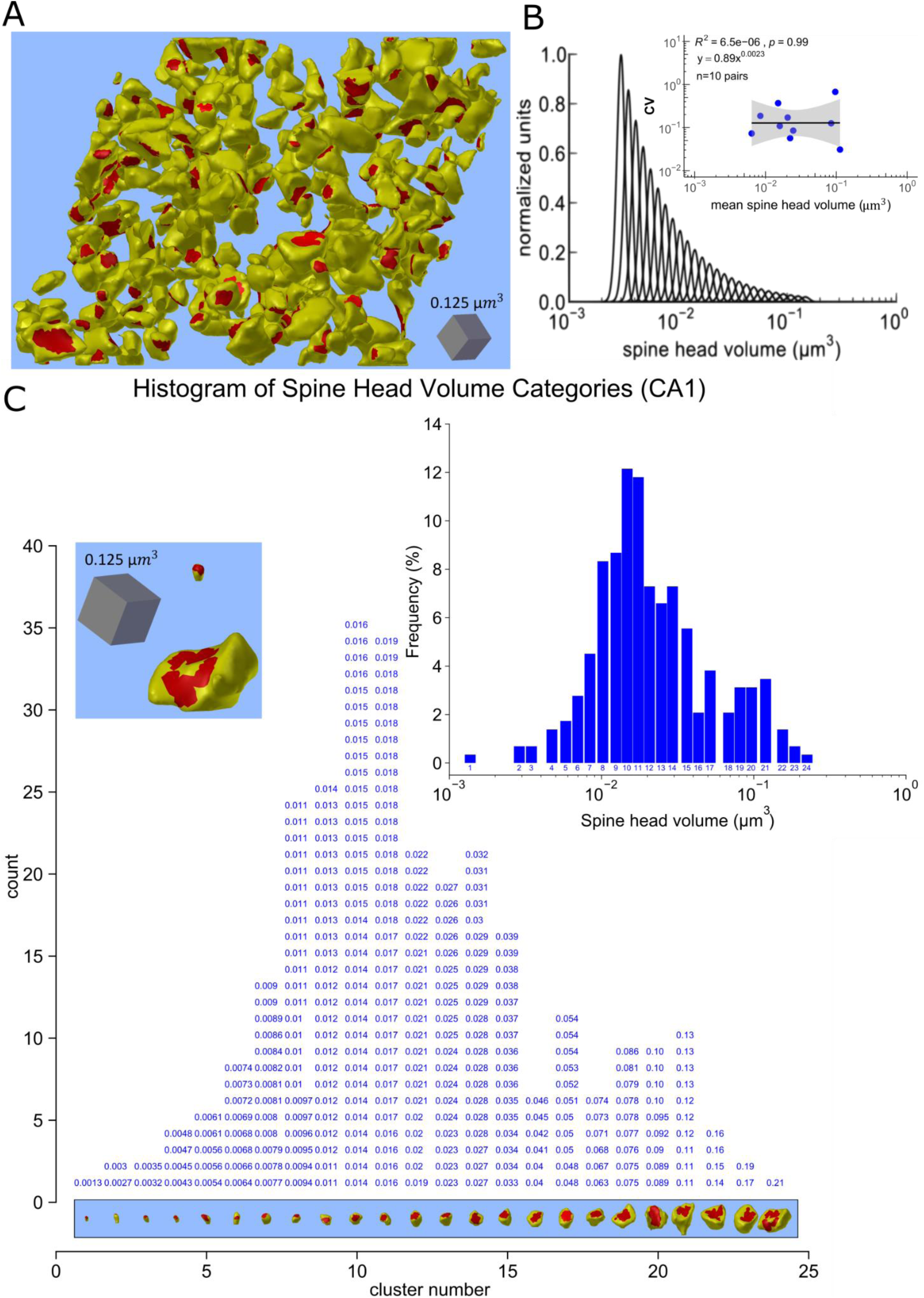
Clustering the spine head volumes from the area CA1 dataset comparing two methods. (A) The 288 spine heads fully captured in the reconstructed volume, displaying the PSD (red) and spine head membrane (yellow). (B) ***Bartol et al., 2015*** using assumptions from signal detection theory showed that 26 distinguishable Gaussian distributions with equal CV (see inset) and overlap of 31% can span the range of spine head volumes of SDSA pairs equivalent to signal to noise ratio of 1 and 69% discrimination threshold common in psychophysics. (C) Our new clustering algorithm (see Algorithm 2, methods) obtained 24 distinguishable categories of all 288 spine heads in the dataset based on the median CV value. The histogram of spine head volumes in log scale is depicted in the panel C inset. The Y axis shows the number of spine head volumes within each category. The actual spine head volumes of the individual spine heads of a given category are stacked vertically in sorted order for that category. The 3D object shown below each category (vertical column) is the actual 3D reconstructed spine head of the largest head volume in the category. The red patches are the postsynaptic densities. The X axis shows the number of distinguishable categories. All spine head volumes are rounded to two significant digits.

The upper left inset contains 3D reconstructions of the smallest and largest spine head volumes. The largest spine in each cluster is illustrated beneath each bin. The highest frequency occurs in cluster #10, which contains 36 spine head volumes (***Fig. 5***C). Interestingly, there appears to be a second peak at around cluster 21.

### Number of Distinguishable States in the Dentate Gyrus MML During Plasticity

It is important to note that under the previous method, as N is increased the scale range factor (defined as the ratio of largest spine head volume to the smallest spine head volume) will always increase as the extremes of the distribution are sampled. This outcome will increase the number of Gaussians that span the range, but will tend to overestimate the true value of N_C_ when the population is not continuous. However, under the new method, as N is increased there will be convergence toward the true value of N_C_ because the true shape of discontinuous distributions are sampled. With the new method we have access to the true frequency of spine head volumes in the clusters, which allows further calculation of the entropy of the distinguishable synaptic strength states, the potential number of modes in the distribution, and the gaps in the range of spine head volumes (bins with no spine head volumes in them).

To explore changes in SISC during synaptic plasticity, we applied the new clustering methods to the four MML datasets from the dentate gyrus (***Fig. 6)***. In order to show the frequency of spine head volumes, the clusters are displayed as histogram bins where each bin is one CV wide. Thus, the CV of spine head volumes within each bin is less than or equal to the median CV found from the SDSA pairs analysis of the specific dataset (see ***Fig. 4***, above). Each bin starts with the smallest spine head volume of the representative cluster and ends with a hypothetical spine head volume that is exactly one CV apart (except for the last bin of each condition that is illustrated with the smallest and largest observed data points in that cluster to better demonstrate the expansion of range of sizes after induction of LTP relative to the controls). As one can observe the spine head volumes enlarged and expanded towards the right hand side of the frequency axis after 30 min and 2 hours post induction of LTP.

**Fig. 6:**
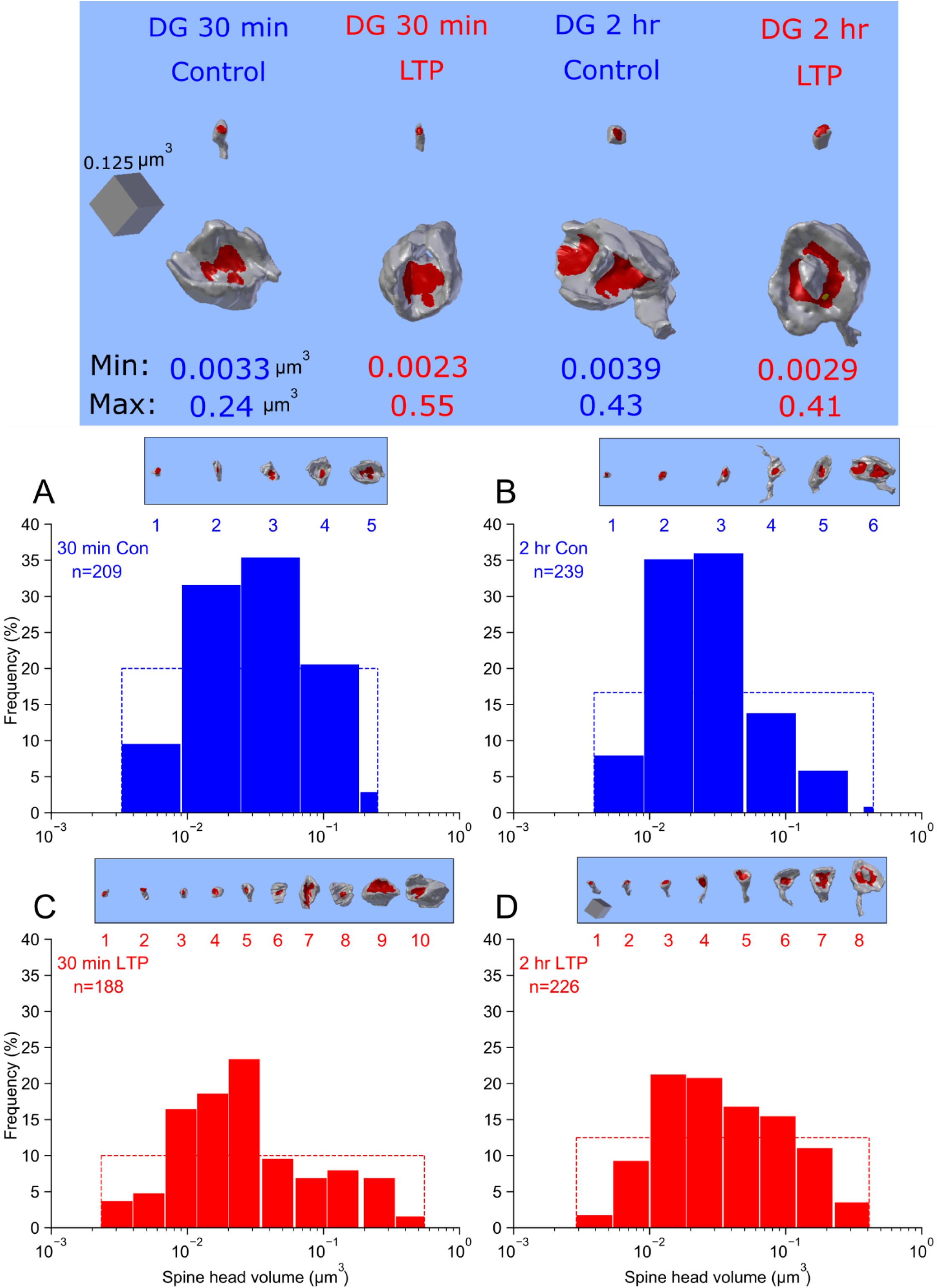
Clustering of spine head volumes in the dentate gyrus datasets. The top panel shows the 3D reconstruction of the smallest and the largest spine head volumes within each dataset with their volumes indicated at two significant digits. Clustering algorithm 2 was used, as in Fig. 1C, to show that following LTP there was an increase in N_C_. Here, categories are illustrated as histogram bins with bin widths equal to the CV shown in Fig. 4 (except that the last bin of each condition is illustrated with observed data points). The categories and actual spine head volume values are shown in **Supplementary Fig. 5-8**. Blue and red colors indicate Control and LTP conditions, respectively. For each panel the Y axis shows the counts of spine head volume in the respective bin divided by total number of spine head volumes in the given dataset and expressed as a percentage. The X axis shows the spine head volumes in *µm*^3^ on a log scale. (A) dentate gyrus 30 min Control, (B) dentate gyrus 2 hr LTP, (C) dentate gyrus 30 min LTP, (D) dentate gyrus 2 hr LTP. The rectangular inset on the top of each histogram shows the largest spine head (on the same scale across panel A-D; scale cube located on panel D inset = 0.125 *µm*^3^) and category number of each category, and aligns with the X axis of the category histogram. For comparison of each histogram to the shape of a uniform distribution, the dashed line indicates the theoretical uniform distribution (with maximum entropy and Shannon information) for the given dataset.

The 30 min control and 2 hr control rats had 5 and 6 distinguishable clusters, respectively. Thus, the values of N_C_ for the control cases were similar despite originating from multiple rats. This closeness between control results validates the repeatability of both the experimental and the computational procedures. At 30 min and 2 hr post-induction of LTP, SISC revealed a higher value of N_C_ due to both the expansion of the scale range factor and to the observable decrease in the CV values (***Fig. 4F; Table 1)***. However, the N_C_ in dentate gyrus either in control or LTP conditions is significantly less than CA1 (N_C_=24). This highly significant difference likely reflects the known differences between area CA1 and dentate gyrus in activation histories and functions in memory formation ***(Snyder, et al., 2001; Saxe, et al., 2006; Krueppel, et al., 2011; Lopez-Rojas, et al., 2016***).

**Table 1:** The number of distinguishable states, or categories, (N_C_) of spine head volumes. For column 3 and 5 the term (± SE), SE stands for standard error calculated using algorithm 1. SRF=scale range factor. The errors on the # of distinguishable clusters is calculated using algorithm 1, 2. For N_C_ Algorithm 1, 2 used with resampling the spine head volumes and using median CV of the observed SDSA pairs for clustering threshold.

| Dataset Type | # SDSA pairs | Median CV of SDSA set | # spine head volumes | Median spine head volumes ($\mu m^3$) | SRF | $N_c$ |
| --- | --- | --- | --- | --- | --- | --- |
| dentate gyrus 30 min Control | 10 | $0.65 \pm 0.12$ | 209 | $0.031 \pm 0.0037$ | 73 | $5 \pm 0.24$ |
| dentate gyrus 30 min LTP | 8 | $0.37 \pm 0.16$ | 188 | $0.022 \pm 0.0014$ | 236 | $10 \pm 0.51$ |
| dentate gyrus 2 hr Control | 18 | $0.56 \pm 0.09$ | 239 | $0.023 \pm 0.0013$ | 110 | $6 \pm 0.48$ |
| dentate gyrus 2 hr LTP | 10 | $0.42 \pm 0.15$ | 226 | $0.031 \pm 0.0036$ | 141 | $8 \pm 0.20$ |
| CA1 | 10 | $0.12 \pm 0.046$ | 288 | $0.018 \pm 0.00091$ | 163 | $24 \pm 0.98$ |

### Shannon Information Storage Capacity of Synapses

The concept of entropy, *H*, comes from the field of thermodynamics and measures the amount of uncertainty or disorder, or the number of possible states of a system. Shannon entropy is defined as the average of Shannon information. Shannon entropy measures the amount of information in the set of distinguishable states, each of which has a probability of occurrence. With more information, it is possible to distinguish more states. The Shannon entropy of a discrete random variable is defined as follows:

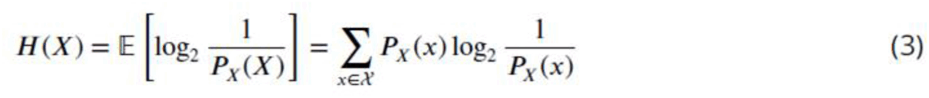

The Shannon information per synapse was calculated from the frequency of spine head volumes *in the distinguishable categories* and using equation (3) where each category is considered a different message. Shannon entropy (bits of information) for the 5 datasets are listed in column four of (Table 2). These data demonstrate that the synapses are not on/off switches and that the induction of LTP increases the information storage capacity of synapses. The maximum number of bits is calculated as the log2(N_C_), which sets an upper bound for SISC. We then compared the spine head volume distinguishable states distributions measured in the control and LTP conditions with a uniform distribution that is maximal entropy among every discrete distribution for a fixed number of states.

**Table 2:** Calculating the entropy of synaptic weights based on the calculated frequency of distinguishable synaptic states. For column 2-5 the term (± SE), SE stands for standard error calculated with bootstrapping using algorithm 1 and 2, concurrently. Spine head volume (SHV). Note that KL(P||U)/*H*(U) is equivalent to KL(P||U)/KL_MAX_.

| Dataset | mean of SHVs | CV of all SHVs | Shannon Entropy $H(P)$ | Maximum Entropy $H(U)$ | KL (P\|\|U) | KL (P\|\|U)/ $H(U)$ |
| --- | --- | --- | --- | --- | --- | --- |
| dentate gyrus 30 min Control | $0.049 \pm 0.0037$ | $1.24 \pm 0.077$ | $2.0 \pm 0.32$ | $2.32 \pm 0.34$ | $0.33 \pm 0.1$ | $0.14 \pm 0.045$ |
| dentate gyrus 30 min LTP | $0.053 \pm 0.0067$ | $1.72 \pm 0.15$ | $3.0 \pm 0.42$ | $3.32 \pm 0.42$ | $0.32 \pm 0.089$ | $0.096 \pm 0.035$ |
| dentate gyrus 2 hr Control | $0.040 \pm 0.0034$ | $1.084 \pm 0.081$ | $2.05 \pm 0.27$ | $2.59 \pm 0.29$ | $0.53 \pm 0.10$ | $0.21 \pm 0.040$ |
| dentate gyrus 2 hr LTP | $0.060 \pm 0.0052$ | $1.41 \pm 0.089$ | $2.74 \pm 0.41$ | $3.0 \pm 0.41$ | $0.26 \pm 0.086$ | $0.086 \pm 0.038$ |
| CA1 | $0.031 \pm 0.0021$ | $1.045 \pm 0.066$ | $4.1 \pm 0.39$ | $4.6 \pm 0.37$ | $0.49 \pm 0.068$ | $0.11 \pm 0.021$ |

**Table 3:** Per animal analysis of SISC. For columns with the term (± SE), SE stands for standard error calculated with bootstrapping using algorithm 1 and 2, concurrently.

| Time points | 30 min |  |  |  | 2 hours |  |  |  |
| --- | --- | --- | --- | --- | --- | --- | --- | --- |
| Animal # | Animal 1 |  | Animal 2 |  | Animal 3 |  | Animal 4 |  |
| Weight (gr) | 623 |  | 448 |  | 648 |  | 490 |  |
| age (postnatal day) | 179 |  | 121 |  | 170 |  | 150 |  |
| time of delta-burst stimulation | 10:30 am |  | 10:30 am |  | 09:30 am |  | 09:30 am |  |
| circadian cycle at delta-burst stimulation | dark |  | dark |  | dark |  | dark |  |
| animal ID | LED50 |  | LED56 |  | LE108 |  | LE113 |  |
| Condition | control | LTP | control | LTP | control | LTP | control | LTP |
| Series code | CLZBJ | TNKPS | NDKZB | MFBCF | JHHZS | HLWLQ | KSGRS | BBCHZ |
| mean fEPSP 5 min pre-delta-burst stimulation % | -3.84 | -3.81 | 3.16 | 2.81 | -2.36 | 0.04 | 0.65 | -2.53 |
| mean fEPSP last 5 min % | -5.38 | 33.61 | 5.28 | 48.33 | -4.76 | 40.68 | -1.10 | 34.45 |
| # SDSA pairs | 6 | 4 | 4 | 4 | 10 | 4 | 8 | 6 |
| SDSA Scale factor | 30 | 7 | 8 | 29 | 26 | 7 | 12 | 107 |
| SDSA Median CV | 0.65 | 0.31 | 0.71 | 0.76 | 0.44 | 0.42 | 0.63 | 0.38 |
| # spine head volumes | 130 | 110 | 79 | 78 | 112 | 111 | 127 | 115 |
| spine head volume Scale factor | 57 | 147 | 64 | 158 | 49 | 110 | 110 | 141 |
| # Clusters for spine head volumes | 5 | 11 | 4 | 5 | 6 | 8 | 5 | 9 |
| Entropy | 1.97 | 3.1 | 1.77 | 1.93 | 2.16 | 2.68 | 1.92 | 2.89 |
| KL divergence | 0.35 | 0.36 | 0.23 | 0.40 | 0.43 | 0.32 | 0.41 | 0.28 |
| # Clusters (Bootstrapping) | 4 ± 0.47 | 10 ± 0.55 | 4 ± 0.39 | 4 ± 0.5 | 6 ± 0.38 | 7 ± 0.71 | 5 ± 0.38 | 8 ± 0.6 |
| Entropy (Bootstrapping) | 1.97 ± 0.06 | 3.1 ± 0.1 | 1.7 ± 0.13 | 1.93 ± 0.1 | 2.16 ± 0.1 | 2.68 ± 0.1 | 1.86 ± 0.1 | 2.89 ± 0.1 |
| KL divergence (Bootstrapping) | 0.14 ± 0.12 | 0.31 ± 0.09 | 0.24 ± 0.12 | 0.27 ± 0.13 | 0.46 ± 0.1 | 0.27 ± 0.1 | 0.42 ± 0.12 | 0.31 ± 0.08 |

### KL Divergence Analysis

Measurement of the distance between an observed distribution, for example spine head volume clusters, and a theoretical uniform distribution with the same number of states is known as the Kullback-Liebler (KL) divergence. A uniform distribution is the maximum entropy discrete probability distribution when there is no constraint on the distribution except having the sum of the probabilities equal 1. Formally, the KL divergence between the distribution of spine head volume clusters (P) and the uniform distribution of states (U) is the difference between cross entropy of (P) and (U) and the entropy of (P): [*H*(*P*, *U*) − *H*(*P*)]. The purpose is to measure the closeness of the empirical probability distribution of the distinguishable states to the maximum entropy distribution with the fixed number of states as the only known constraint on the distribution. When the distribution of the distinguishable synaptic states is close to uniform, the KL divergence will be low and the Shannon entropy will be nearly maximized. Here we are measuring the fundamental limits of SISC and not the likely efficient coding occurring at the population level with the more uniform distinguishable bins. The spine head volume distribution in CA1 is compared with the uniform distribution with 24 states in **Supplementary Fig. 4** and quantified in Table 2.

The maximum entropy and KL divergence were all quite low for all four datasets from the dentate gyrus and were around 50% lower than for CA1 (Table 2, column 6; **Supplementary Fig. 4**). The KL divergence for the 30 minute LTP was not different from the control, but the KL divergence for the 2 hour LTP was less than half of its matched control condition (Table 2, column 4). This means that the changes in the distribution of spine head volumes that occurred between 30 minutes and 2 hours shifted toward a more nearly uniform distribution having maximum information due to optimal use of the distinguishable states.

Moreover, we calculated the ratio of the KL divergence values over the maximum value that KL divergence can possibly have in each case (KL / KL_MAX_) where KL_MAX_ equals *H*(*U*). The purpose is to measure the ratio of the measured KL divergence value to the maximum value it could hypothetically have (Table 2, column 7). The lower the ratio, the more efficient is the usage of distinguishable states for the storage of synaptic strength values across the population of synapses. There was a 31% increase in efficiency 30 minutes following LTP and an additional 17% increase after 2 hours. These findings imply that LTP moves distributions of spine head volumes toward maximum information storage limit and maximum efficiency due to optimal use of the distinguishable states.

## Discussion

This paper introduces a new analytical approach for determining SISC that has several advantages over our prior approach (***Bartol et al., 2015***). The new method was used on data from area CA1 to compare it with the prior approach. It was then applied to track SISC changes in the dentate gyrus at 30 minutes and 2 hours following the induction of LTP. The analyses revealed that synaptic precision, based on covariance of spine head volume in SDSA pairs, was not altered during LTP. This finding suggests that as one spine of the pair enlarged (or shrank) following LTP, the other spine head changed in tandem. Note that SDSA pairs are independent synapses, but because they are driven by the same input they respond in a similar way and arrive at the same size due to the precision of the underlying mechanisms they each possess. We also found that the number of distinguishable synaptic strengths is increased during LTP by altering the range and frequency of spine head volumes in the distinguishable clusters. At 30 minutes following induction of LTP, spine head volumes shifted from the middle of the range both towards smaller and larger sizes. Thus, consistent with findings in CA1 in rat in ***Bourne and Harris, 2011*** DBS LTP induction triggered spinogenesis followed by loss of small excitatory synapses and a subsequent enlargement of the remaining synapses by 2 hr. These data suggest that dendritic segments of granule cells in the dentate gyrus coordinate their structural plasticity. Coordination occurs across the complete range of synapses and maintains a homeostatic balance of excitatory inputs. The mechanism for this coordination could involve local protein synthesis and selective capture or redistribution of dendritic resources, as were observed in CA1 in rat in ***Bourne and Harris, 2011***.

We observed an increase in the number of bits from 2.0 bits in the control conditions for both time points to 3.0 bits after 30 min and 2.7 bits at 2 hr following induction of LTP. These outcomes were a consequence of two changes to the distribution of spine head sizes. First, there was an increase in both the larger and smaller spine head volumes, which broadened the size distributions. In addition, there was a change in the frequency of spine head volumes in each bin resulting in a more uniform distribution of spine head volumes. These changes brought the overall distribution of spine head volumes (***Fig. 6*** C and D) closer to the optimal distribution with maximum Shannon information (Table 2 column 5). This broadening in the size range was observed in both the 30 min and the 2 hr difference distributions. As a consequence, the information storage capacity was increased by around 50% following the LTP induction, an increase that was preserved between 30 min and 2 hr.

Moreover, there was also evidence for further reorganization of spine head volumes after 30 min. The KL divergence is a measure of the difference between two distributions; when applied to the distributions of spine head volumes and the maximal entropy uniform distribution, the value at 2 hr after LTP induction was half of that of 30 min following LTP induction relative to their controls, thereby using the range of spine head volume sizes more efficiently. This analysis suggests that after learning has taken place, underlying mechanisms continue to push the distribution of spine head volumes toward a more efficient storage of information at the population level.

### Advantages of the new SISC analysis

There are several advantages to the new SISC method for assessing the storage capacity of synapses. Signal detection theory (***Bartol et. al, 2015***) assumed that the width of the Gaussian curves, based on the CV of the SDSA pairs, were distributed equally along the full range of sampled spine head volumes, without accounting for gaps in the distribution. Thus, the signal detection theoretical approach tended to overestimate the true number of distinct synaptic states because the distribution of the population was assumed to be continuous. In the new SISC analysis, the number of distinct synaptic states defined by the individual clusters converges toward the true number of states as the number of spine head volumes increases and the true shape of the distribution is sampled. A second advantage is that the full population of spine head volumes, not just the SDSA pairs, are included in the analysis, greatly improving the statistical power of the estimate. A third advantage is that there are no free parameters in the estimate, unlike signal detection theory where the degree of overlap of the Gaussians is a free parameter. A fourth advantage is that the new method is robust to the outliers. The largest spine head volume we found in an earlier data set (***Harris and Stevens, 1989***) in rat hippocampal area CA1 was 0.55 cubic microns. It is worth mentioning that by adding this one value to the 288 CA1 spine head volumes would result in 25 distinguishable clusters, an increment of 1 state using SISC. Using the prior signal detection approach results in 39 distinguishable Gaussians spanning the range, a 50% increase in the number of states. The previous method is not robust to outliers. Finally, the new method, based on information theory, allows access to the frequency of the clusters, making it possible to compute the entropy of the distinguishable synaptic strengths, the number of modes in the distribution, and any gaps in the range of functionally distinguishable synaptic strengths.

Our approach can be applied broadly to cluster these and other measures of synaptic efficacy in different brain regions and model organisms. Furthermore, the impact of other measures of synaptic plasticity can now be assessed reliably, such as changes in spine neck dimensions and pre- or postsynaptic dimensions and composition. In addition, it will be possible to determine how changes in the location or dimensions of key subcellular components, such as mitochondria, smooth endoplasmic reticulum, polyribosomes, endosomes, and perisynaptic astroglia, affect SISC. These opportunities to generalize SISC will surely result in greater understanding about the role of each component individually, and in concert, in determining normal synaptic weight and plasticity. Ultimately, the outcomes should give insight into how disrupted synapses result in cognitive decline during aging or as a consequence of neurological disorders.

### Information Theory of Synapses

Information theory has been applied to analysis of spike trains ***(Dayan, P. and Abbott, L.F., 2005)*** but has not been used at the level of synaptic strength. We have shown that the amount of information represented by synaptic weights in neural circuits can be quantified by the distinguishability of synaptic weights. Here “distinguishability” fundamentally depends on the precision of the synaptic weights. When the precision of synaptic weights is low, the amount of information that can be stored in the ensemble of the neurons will also be low. Complete absence of precision implies a random process for setting synaptic weight and no information being stored at synapses. Because spine head volume is highly correlated with synapse size (***Bartol et al., 2015***), the precision of spine head volumes can be used to measure the distinguishability of the synaptic weights. High precision yields a greater number of distinguishable spine head volume clusters and hence higher information storage capacity. The number of distinguishable weights is not static but varies with the history of synaptic plasticity and is different in different parts of the brain. Thus, the amount of information that a population of synapses can store is not fixed but can be changed.

We made comparisons with the uniform distribution because it is the most conservative assumption when biological constraints (e.g. fixed values for the shape, mean, median, max, …) on the spine head volume distribution are not known. In our mathematical setting when no constraint is applied (except the number of states being fixed) the uniform distribution has maximal entropy among every discrete distribution. This is why the uniform distribution for a fixed number of states is a “lower bound” on optimality and an upper bound for SISC. For example, we measured the entropy of the observed synaptic weight distribution in area CA1 based on the probability of the 24 distinguishable states as having a value of 4.1 bits while the maximal entropy distribution when having 24 states is the uniform distribution with 4.59 bits of information. We used KL divergence analysis to measure the closeness of the spine head volume distributions to the maximum entropy distribution. What is remarkable here is while LTP does not change the number of spines, axons, or the summed synaptic area per unbiased length of dendrite in the dentate gyrus, the late phase of LTP pushes the distribution of distinguishable synaptic states closer to the upper bound of SISC given by the maximal entropy distribution.

If biological constraints were found at some point in the future, we could calculate the maximal entropy distribution under those constraints and measure the distance between the empirical distribution entropy (e.g., log normal) with the aforementioned maximal entropy distribution (with optimized parameters under those constraints). We can speculate that under those conditions the KL divergence would be even smaller. Because no biological constraint is known, we chose the universal maximal entropy discrete distribution as a reference frame to compare with and check optimality.

It is worth noting that synaptic activity and hence SISC are highly variable and change with an animal’s behavior. This manuscript is only concerned with the optimality of information storage capacity based on the synaptic weight itself as the unit of storage of information. The question of how optimally synaptic activity is used to store working memories in neural circuits and how efficiently the synaptic weight is used in those codes is beyond the scope of this manuscript and is left for future research.

### Mechanisms underlying the SISC Increase

Observations of increases and decreases in strength of synaptic potentials following LTP were determined using electrophysiology measurements. Our SISC analysis suggests that the changes in strength are due to increases in larger spines and smaller spine head sizes, which is consistent with observations based on light microscopy. In addition, we have observed decrease in CV among SDSA pairs (i.e., higher precision).

It should be noted that the increase in SISC via change in spine size has two principal contributing mechanisms rooted in broadening the range of spine head volumes. One is the increase in the largest spine head volumes due to LTP. But an equally important change is the concurrent and offsetting reduction in spine sizes in other synapses. In area CA1, a homeostatic mechanism was revealed where induction of LTP with saturating theta burst stimulation enlarged some synapses by 2 hr post induction, while stalling the outgrowth of small dendritic spines (***Bourne and Harris, 2011; Bell et al., 2014***). This effect resulted in a stable total summed synaptic weight per unit length of dendrite. Furthermore, the enlarged spine synapses also stabilized their neighboring smaller spines into efficient synaptic clusters ***(Chrillo et al., 2019)***. Thus, future analysis may reveal zones with high SISC and others with low SISC even along short segments of dendrite.

Here we further elaborate on the effect of LTD on the broadening of sizes. Particularly, concurrently induced “heterosynaptic” LTD has long been known to occur in the dentate gyrus perforant pathways (***Abraham & Goddard, 1983***) including after *in vivo* 400 Hz DBS as used in the current study (***Bowden et al., 2012***). We have shown in earlier works that this form of LTD is in fact activity-dependent, implying a reduction in the threshold such that either constitutive or evoked synaptic activity in the non-tetanized synapses becomes transiently capable of evoking the LTD (***Abraham et al., 2007***). ***Jedlicka et al. (2015)*** showed in a biophysically realistic compartmental granule cell model that this pattern of results can be accounted for by a voltage-based spike-timing-dependent plasticity (STDP) rule combined with a relatively fast Bienenstock-Cooper-Munro (BCM)-like homeostatic metaplasticity rule, all on a background of ongoing spontaneous activity in the input fibers. These results suggest that, at least for dentate granule cells, the interplay of STDP and BCM plasticity rules and ongoing pre- and postsynaptic background activity determines not only the degree of input-specific LTP elicited by various plasticity-inducing protocols, but also the degree of “associated LTD” in neighboring non-tetanized inputs, as generated by the ongoing constitutive activity at these synapses.

Furthermore, ***Yang et al. (2008)***, after simultaneously monitoring EPSPs and dendritic spines using combined patch-clamp recording and two-photon time-lapse imaging in the same CA1 pyramidal neurons in acute hippocampal slices, showed that the initial expression of LTP and spine expansion are dissociable, but that there is a high degree of mechanistic overlap between the stabilization of structural plasticity and LTP. In our results, we showed that the coefficient of variation of spine head volumes (of the complete sample) of DG 30 min LTP (1.72) increased ∼40% in comparison to DG 30 min Control (1.24) showing the extent of the expansion of spine head volumes. The CV value rested at 1.41 for the DG 2 hr LTP which is ∼30 % higher than the DG 2hr Control (1.084).

These mechanisms thus could explain much of the broadening of the range of sizes which by itself would increase SISC. However, we also observed a consistent though non-significant decrease in CV of SDSA pairs, and this is a third mechanism that could contribute to the increase in SISC. As yet we do not have an explanation for the decrease in CV, and this remains a focus for future research. However, the strong DBS protocol for inducing LTP may have served to increase the coordinated activation histories of the SDSA pairs, leading to the CV reduction.

### Comparison to Synapses in the Cerebral Cortex

We observed a low median CV (∼0.1) among SDSA pairs in CA1 and a larger median CV (∼0.6) in DG with no significant trend with spine head volume, but a large (∼1 order of magnitude) variation from pair to pair. Thus, our results reveal that CV among SDSA pairs differs across brain regions.

SDSA pairs are a type of “joint synapse,” but note that joint synapses have a broader definition — namely, *multiple synapses* sharing the same pre- and postsynaptic *neurons*, not just the same axon and the same dendrite. Joint synapses with up to 7 shared synapses across the entire neuron have been found in other 3DEM studies (Dorkenwald et al., 2019), (Motta et al., 2019). A direct comparison of our study to these other studies is difficult to make due to multiple differences in experimental design. There are at least 8 major differences between our study and that of Motta et al. (2019): 1) choice of model organism (mouse vs. rat); 2) brain region (somatosensory cortex of mouse vs. CA1 and DG of rat hippocampus); 3) measurement criterion (ASI vs. spine head volume); 4) connection type (joint synapses in general vs. SDSA pairs specifically); 5) sample size (thousands vs. tens); 6) measurement conditions (control only vs. control and LTP); 7) number of animals (1 vs. 3 in CA1 and 4 in DG); 8) analysis of measurement error (none vs. error estimation based on 4 independent trials). There are a number of differences in how anatomical data was analyzed from the hippocampus and the cortex. First, the surface area of the axon-spine interface (ASI), not spine head volume, was measured in layer 4 of the somatosensory cortex (***Motta et al., 2019***). Motta et al. noted that saturating LTP or LTD could explain the lower CV (higher level of precision) which they observed among *joint synapses* with the largest or smallest ASI. Second, ***Motta et al. (2019)*** also observed higher CV among spines with intermediate sized ASI, which is inconsistent with our findings in ***Fig. 4***. Perhaps this difference can be attributed to the possibility that ASI depends on aspects of synaptic function other than synaptic weight. Third, it is critical to point out that the measurement error needs to be smaller than the estimated synaptic plasticity precision (CV). ***Motta et al. (2019)*** neither explored the precision of synaptic plasticity nor the measurement error of ASI. Comparison of their study with ours will require further analysis of their data. Fourth, another important difference between the two studies is the experimental condition of the animal. ***Motta et al. (2019)*** stated in their abstract, “We quantified connectomic imprints consistent with Hebbian synaptic weight adaptation, which yielded upper bounds for the fraction of the circuit consistent with saturated long-term potentiation,” assuming that synapses that had undergone saturated LTP were the synapses with low CV. However, the CVs of synapses in our CA1 data were equally precise for small spines as for large spines. Finally, comparison of our results in CA1 and DG revealed that: 1) different brain regions have different levels of precision and 2) within a region the precision level varies among SDSA pairs by over an order of magnitude within control and 30 min and 2 hr post LTP induction datasets (Fig. 4), but with no significant trend from the smallest to the largest spine head volume.

In another study of pyramidal neurons in cortical layer 2/3 (***Dorkenwald et al., 2019***), spine head volumes were similar among pairs that shared the same axon but were on different dendrites from the same cell. The distribution of spine head volumes in their sample had two broad peaks, suggesting that the populations of small and large synapses were distinct. Similarly, despite the small numbers, there may be two distinct peaks in our distribution of spine head volumes in area CA1. The frequency of small spines is much higher than larger spines. And small spines are generally more transient than the larger, typically more stable spines ***(Holtmaat et al., 2009)***. Indeed, in adult hippocampal area CA1, small spine outgrowth is stalled while synapses on the largest stable spines further enlarge following LTP (***Bell et al., 2014***), which is consistent with our observation of an increase in the frequency of large spines 2 hours after induction of LTP (***Fig. 2F***).

## Conclusion

This paper explored the precision of synaptic plasticity and synaptic information storage capacity. Our methods and set of algorithms revealed new insights about information storage capacity at synaptic resolution. Information storage and coding in neural circuits has multiple substrates over a wide range of spatial and temporal scales. How information is coded, and its potential efficiency depends on how the information is stored. The synaptic weight is itself the information that is stored, and this information is retrieved by synaptic strength as assayed with test pulses. Here we measured the efficiency of information storage by analyzing the number of distinguishable categories for synaptic weights and the rearrangement of the spine sizes following LTP induction. The interplay of LTP and LTD plasticity rules after induction of our DBS protocol broadened the spine size range and reduced the CV. Based on comparing the distributions of synaptic weights and that of a uniform weight distribution (the maximum entropy distribution), we found that LTP is highly efficient in storing information in synaptic weights across the distinguishable clusters. The weights continued to evolve over time towards a maximal entropy distribution, thereby optimally using the number of distinguishable clusters. From the perspective of information theory, this analysis has revealed a new way that the late phase of LTP may be involved in shifting the distribution of spine head volumes to achieve a more efficient use of coding space in the available synaptic population.

## Methods and Materials

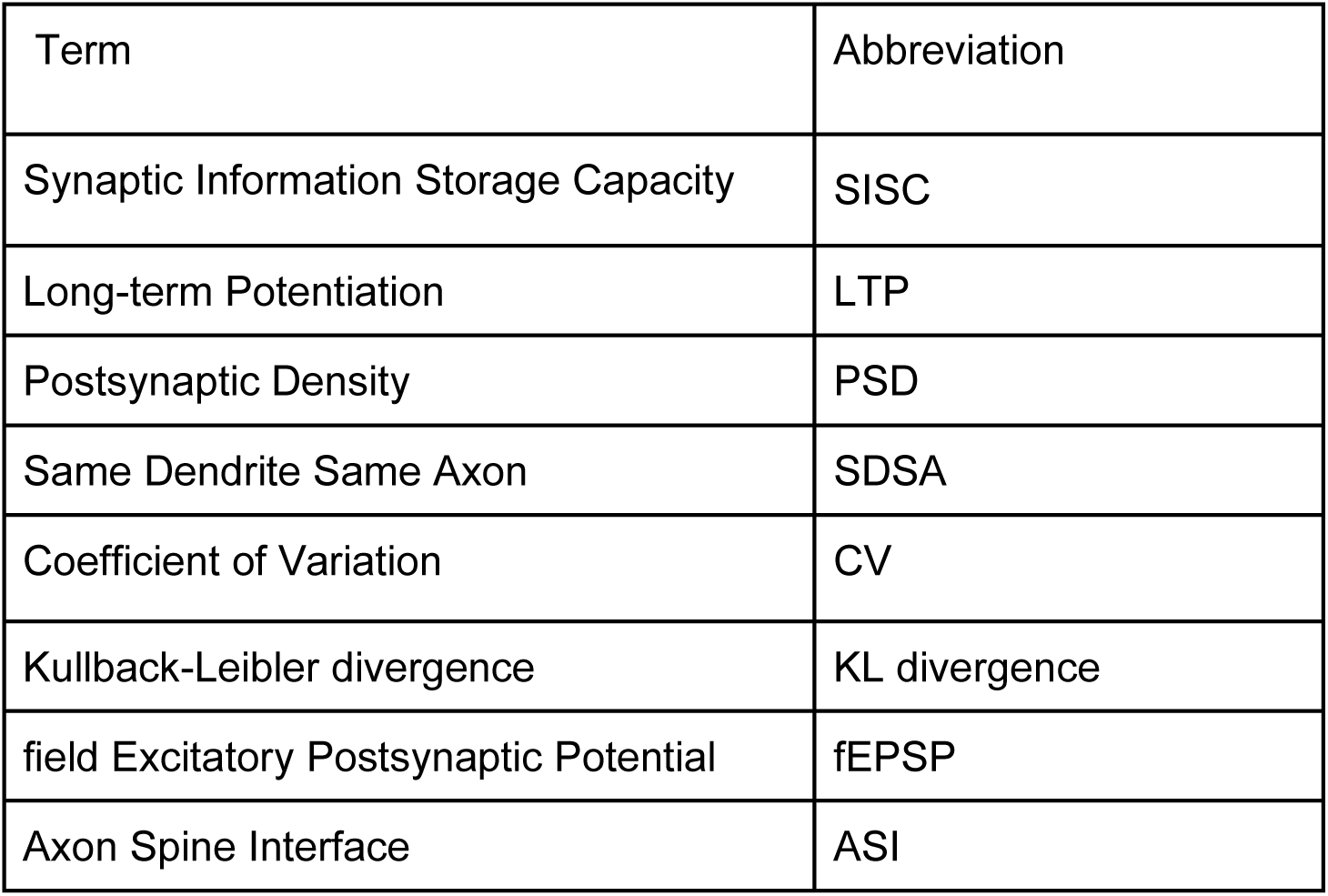

| Term | Abbreviation |
| --- | --- |
| Synaptic Information Storage Capacity | SISC |
| Long-term Potentiation | LTP |
| Postsynaptic Density | PSD |
| Same Dendrite Same Axon | SDSA |
| Coefficient of Variation | CV |
| Kullback-Leibler divergence | KL divergence |
| field Excitatory Postsynaptic Potential | fEPSP |
| Axon Spine Interface | ASI |

For the experiments done for the CA1 dataset, see the materials and methods section in ***Bartol et al., 2015***. Here we present the details of experiments done for the 30 min and 2 hours post induction of LTP in 4 rats. Explanations are the modified version of 30 min control and LTP datasets published in ***Bromer et al., 2018***.

## Surgery and Electrophysiology (dentate gyrus)

The 30 min dataset was collected from two adult male Long-Evans rats aged 121 and 179 d at the time of LTP induction and perfusion. The 2 hour dataset was collected from two adult male Long-Evans rats aged 150 and 170 d at the time of LTP induction and perfusion. It is worth noting that the histograms in ***Fig. 2***A,D and ***Fig. 2***B,E are made for the combined datasets from the two 30 min datasets for control and LTP conditions, and the two 2 hour datasets for control and LTP conditions, respectively.

The animals were surgically implanted using stereotaxic coordinates as previously described in ***Bowden et al., 2012***[CA1] with wire stimulating electrodes placed separately into the medial and lateral perforant pathways running in the angular bundle in the LTP hemisphere, and in only the medial perforant pathway in the control hemisphere due to the limited number of channels in the animal’s head-plug connector and our primary interest in the medial path data which are described in this paper. Wire field excitatory postsynaptic potential (fEPSP) recording electrodes were implanted bilaterally in the dentate hilus. Two weeks after surgery, baseline recording sessions (30 min and 2 hours) commenced, with animals being in a quiet alert state during the animals’ dark cycle. Test pulse stimuli were administered to each pathway as constant-current biphasic square-wave pulses (150 *µs* half-wave duration) at a rate of 1/30 s, and alternating between the three stimulating electrodes. The test pulse stimulation intensity was set to evoke medial path waveforms with fEPSP slopes > 3.5 mV/ms in association with population spike amplitudes between 2 and 4 mV, at a stimulation current ≤ 500 *µA*. Paired-pulse and convergence tests were used to confirm successful stimulating placements in the MPP and LPP, as previously described in ***Bowden et al., 2012***. On the day of LTP induction, after stable baseline recordings were achieved, animals received 30 min of test pulses followed by delta-burst stimulation delivered to the ipsilateral medial perforant path, while the contralateral hippocampus served as a control. The LTP-inducing delta-burst stimulation protocol consisted of five trains of 10 pulses (250 *µs* half-wave duration at the same pulse amplitude as for the test pulses) delivered at 400 Hz at a 1 Hz interburst frequency, repeated 10 times at 1 min intervals (***Bowden et al., 2012***). Test pulse stimulation then resumed until the brains were obtained at 30 min and 2 hours after the onset of delta-burst stimulation. The initial slope of the medial path fEPSP was measured for each waveform and expressed as a percentage of the average response during the last 15 min of recording before delta-burst stimulation.

## Perfusion and Fixation (dentate gyrus)

At 30 min or 2 hr after the commencement of delta-burst stimulation, animals were perfusion-fixed under halothane anesthesia and tracheal supply of oxygen (***Kuwajima et al., 2013***). The perfusion involved brief (∼20 s) wash with oxygenated Krebs-Ringer Carbicarb buffer [concentration (in mM): 2.0 CaCl2, 11.0 D-glucose, 4.7 KCl, 4.0 MgSO_4_, 118 NaCl, 12.5 Na_2_CO_3_, 12.5 NaHCO_3_; pH 7.4; osmolality, 300–330 mmol/kg], followed by fixative containing 2.0% formaldehyde, 2.5% glutaraldehyde (both aldehydes from Ladd Research), 2 mM CaCl_2_, and 4 mM MgSO_4_ in 0.1 M cacodylate buffer (pH 7.4) for ∼1 hr (∼1,900 mL of fixative was used per animal). The brains were removed from the skull at about 1 hr after end of perfusion, wrapped in several layers of cotton gauze, and shipped on ice in the same fixative from the Abraham Laboratory in Dunedin, New Zealand, to the laboratory of K.M.H. in Austin, Texas by overnight delivery (TNT Holdings B.V.).

### Tissue Processing and Serial Sectioning (dentate gyrus)

The fixed tissue was then cut into parasagittal slices (70 µm thickness) with a vibrating blade microtome (Leica Microsystems) and processed for electron microscopy as described previously (***Kuwajima et al., 2013***), (***Harris et al., 2006***). Briefly, the tissue was treated with reduced osmium (1% osmium tetroxide and 1.5% potassium ferrocyanide in 0.1 M cacodylate buffer) followed by microwave-assisted incubation in 1% osmium tetroxide under vacuum. Then the tissue underwent microwave-assisted dehydration and en bloc staining with 1% uranyl acetate in ascending concentrations of ethanol. The tissue was embedded into LX-112 epoxy resin (Ladd Research) at 60° C for 48 hr before being cut into series of ultrathin sections at the nominal thickness of 45 nm with a 35° diamond knife (DiATOME) on an ultramicrotome (Leica Microsystems). The serial ultrathin sections from MML (region of molecular layer ∼125 µm from top of granule cell layer in dorsal blade of the hippocampal dentate gyrus) were collected onto Synaptek Be-Cu slot grids (Electron Microscopy Sciences or Ted Pella), coated with Pioloform (Ted Pella), and stained with a saturated aqueous solution of uranyl acetate followed by lead citrate (***Reynolds, 1963***).

### Imaging and Alignment (dentate gyrus)

“The serial ultrathin sections were imaged, blind as to condition, with either a JEOL JEM-1230 TEM or a transmission-mode scanning EM (tSEM) (Zeiss SUPRA 40 field-emission SEM with a retractable multimode transmitted electron detector and ATLAS package for large-field image acquisition; (**Kuwajima *et al., 2013***)). On the TEM, sections were imaged in two-fleld mosaics at 5,000× magnification with a Gatan UltraScan 4000 CCD camera (4,080 pixels × 4,080 pixels), controlled by DigitalMicrograph software (Gatan). Mosaics were then stitched with the photomerge function in Adobe Photoshop. The serial TEM images were first manually aligned in Reconstruct (***Fiala JC., 2005***; synapseweb.clm.utexas.edu/software-0) and later in Fiji with the TrakEM2 plugin (refs. (***Fiala JC., 2005; Cardona A, et al., 2012; Schindelin J, et al., 2012***); flji.sc). On the tSEM, each section was imaged with the transmitted electron detector from a single field encompassing 32.768 µm × 32.768 µm (16,384 pixels × 16,384 pixels at 2 nm/pixel resolution). The scan beam was set for a dwell time of 1.25–1.4 ms, with the accelerating voltage of 28 kV in high-current mode. Serial tSEM images were aligned automatically using Fiji with the TrakEM2 plugin. The images were aligned rigidly first, followed by application of affine and then elastic alignment. Images from a series were given a five letter code to mask the identity of experimental conditions in subsequent analyses with Reconstruct. Pixel size was calibrated for each series using the grating replica image that was acquired along with serial sections. The section thickness was estimated using the cylindrical diameters method (***Fiala et al., 2001***).” (***Bromer et al., 2018***))

### Unbiased Reconstructions and Identfication of SDSA Pairs (dentate gyrus)

Three dendrites of similar caliber were traced through serial sections from each of the two control and two LTP hemispheres for a total of six dendrites per condition (with a total of 24 dendrites for the 30 min and 2 hr datasets). Dendrite caliber previously has been shown to scale with dendrite cross-sectional area and microtubule count (***Harris et. al., 2022***). The microtubule count ranged from 30 to 35 and represents the average among all dendrites found in the MML of dentate gyrus. These dendritic segments ranged in length from 8.6 to 10.6 µm for the six control dendrites and 9.3 to 10.6 *µm* for the six LTP dendrites. All spines were identified visually by flickering between two images from the adjacent serial sections, to detect small changes in membrane curvature at the spine origins. This was repeated along the entire length of each sampled dendrite in the series.

Once identified, the spines were segmented by manually drawing closed contours along the plasma membrane (including spine head and neck) in each of the aligned images they appear. The spine contours were laid down to remove them at their origins from the dendritic shaft. The spine origin appears as an inflexion of the plasma membrane at the junction between spine neck and dendritic shaft, and this was visually confirmed both in 3D reconstructions and the underlying serial EM images.

In each spine, the PSD was identified by its electron density along the postsynaptic plasma membrane and the presence of small synaptic vesicles accumulating in the opposing presynaptic bouton. Closed contours were created to encapsulate the PSD and the presynaptic active zone. These contours crossed the postsynaptic plasma membrane perpendicularly to ensure that the correct area of the membrane surface mesh could be assigned as the PSD later in Blender (see below).

Both spine and PSD traces were curated by experts to ensure their accuracy. All manual segmentation and trace curation were done by experienced tracers who remained blind as to the experimental conditions (i.e., stimulation and layer location). A total of 209 spines were complete along the control dendrites and 188 spines were complete along the LTP dendrites. These were used for the indicated analyses. The unbiased dendritic segment analysis involved assessing the number of synapses, SDSAs, and axons interacting with each dendritic segment. Beginning in the center of each of the 24 dendrites, the presynaptic axons were traced past the nearest neighboring axonal bouton until they were determined to form synapses with the same dendrite or a different dendrite. Only the middle portion of the dendrite lengths could be used because only spines in the middle of the dendrite had presynaptic axons sufficiently complete within the series to determine their connectivity. In three cases, one axon made synapses with dendritic spines from two different dendrites in our sample, and these three were included for both dendritic segments. Each of the 24 dendrites (12 for 30 min and 12 for the 2 hr datasets) was truncated to contain the central 15-20 spine and shaft synapses with known connectivity. The z-trace tool in Reconstruct was used to obtain the unbiased lengths spanning the origin of the first included spine to the origin of the first excluded spine ***(Fiala et al., 2001)***. The lengths ranged from 2.8 to 5.9 *µm* for the six control dendrites and 3.1 to 6.1 *µm* for the six LTP dendrites. Then the number per micrometer length of dendrite was computed for spines, axons, and SDSAs. PSD areas were measured in Reconstruct according to the orientation in which they were sectioned ***(Harris et al., 2015)***. Perfectly cross-sectioned synapses had distinct presynaptic and postsynaptic membranes and synaptic cleft, and their areas were calculated by summing the product of PSD length and section thickness for each section spanned. En face synapses were cut parallel to the PSD surface, appeared in one section, and were measured as the enclosed area on that section. Obliquely sectioned PSDs were measured as the sum of the total cross-sectioned areas and total en face areas without overlap on adjacent sections. Then the synapse areas were summed along the truncated, unbiased dendritic length to compute values.

### Segmentation and Evaluation of Spines (dentate gyrus)

“Blender, a free, open-source, user-extensible computer graphics tool, was used in conjunction with 3D models generated in Reconstruct. We enhanced our Python add-on to Blender, Neuropil Tools (***Bartol et al., 2015***), with a new Processor Tool to facilitate the processing of the 3D reconstruction and evaluation of spines. The additions encompassed in Processor Tool were as follows:

i. The software allows for the selection of traced objects from Reconstruct (.ser) files by filter, allowing the user to select only desired contour traces (in this case spine head and PSD contours for three dendrites per series).
ii. At the press of a button, the tool generates 3D representations of selected contours in Blender. This step invokes functions from VolRoverN (***Edwards et al., 2014***) from within Blender to generate mesh objects by the addition of triangle faces between contour traces.
iii. Smoothing and evening of the surface of spine objects is accomplished with GAMer software (fetk.org/codes/gamer/).
iv. In a few cases, the formation of triangles was uneven and required additional manipulation by Blender tools and repeating of step iii before proceeding to step v.
v. Finally, the PSD areas are assigned as metadata (represented by red triangles) for the reconstructed spine heads.

This assignment is performed based on the overlap of PSD and spine head contours (described above) in 3D space. Dendritic spines were segmented as previously described (***Bartol et al., 2015***) using the Neuropil Tools analyzer tool. We chose to measure spine volumes because at present they can be more accurately measured than other correlated metrics, synaptic area and vesicle number (***Bartol et al., 2015***). The edges of the synaptic contact areas are less precisely determined in oblique sections, and vesicles can be buried within the depth of a section or span two sections and hence are less reliably scored. The selection of spine head from spine neck and from spine neck to dendritic shaft were made using the same standardized criterion as before (visually identified as halfway along the concave arc as the head narrows to form the neck). Spines were excluded if they were clipped by the edge of the image dataset. To ensure the accuracy of the measurements, segmentation of the spine head volume was performed four times (twice each by two people) and averaged. A further check was added at this step, whereby spine heads with a CV ≥ 0.02 for all four measurements were visually evaluated by an expert, and any discrepancy in the segmentation was corrected. Interestingly, the only spines with a CV larger than 0.02 were in the LTP condition. We believe this occurs because the spines undergoing LTP are likely to be in transition at the 30 min time point, and as such the delineation between spine head and spine neck is more difficult to distinguish. In the two control condition series, further evaluation by an expert was performed, and adjustments were made accordingly (***Bromer et al., 2018***))” (Fig. 2 and Fig. S4 in (***Bromer et al., 2018***)).

## Code Availability

The data and codes used in the present study will be available in the following github link: https://github.com/MohammadSamavat

### Statistical Analysis

Statistical analysis and plots were generated using Python 3.4 with NumPy, SciPy, and Matplotlib. ***Fig. 4*** is made by R programming packages as follows: ggplot2, ggpubr, scales, xlsx, ggpmisc.

In order to show the empirical distribution of spine head volumes for the 4 dentate gyrus datasets, we illustrated the 4 dentate gyrus spine head volume histograms in ***Fig. 2***. For panel A-D the Y axis shows the frequency of spine head volumes within each of the bins and the X axis shows the bins start and end points. To get the bins’ start and end points, the range of the 4 datasets were divided into 11 equal width bins (identical bins for all 4 dentate gyrus datasets) in logarithmic scale (Fig. 2 panel A-D).

We designed a statistical test based on nonparametric bootstrap hypothesis testing to see if the observed peaks and troughs of the trajectories in ***Fig. 2***E,F are significant. To generate each of the 10,000 bootstrap samples we resampled the combined 4 DG dataset (n=862) with replacement and extracted DG 30 min control (n=209) and LTP (n=188) along with DG 2 hr control (n=239) and LTP (n=226). For each of the bootstrap samples we calculated the test statistic (summation of differences of frequencies) for all peaks and troughs. The p-value is calculated as the ratio of number of cases that the test statistics was as extreme as that of observed peaks and troughs of the empirical data over 10,000. The list of test statistics and p-values are shown in the following table.

**Supplemental Table 1:** P value of the observed peaks and troughs in Fig 2 E,F. A=freq_DG_30min_Control, B=freq_DG_30min_LTP, C=freq_DG_2hr_LTP, D=freq_DG_2hr_Control. e.g. sum(D - B): It means we have first subtracted the frequencies of the 30 min LTP histogram from that of 2 hr LTP for each represented bin and then summed the differences (the other statistics are defined in the same way). Note: the P value 0 in row 3 means in the 10,000 bootstrap samples there were no cases with summation of differences being as extreme as the observed value.

| statistics | Observed value<br>(for SHV<median) | P-Value | Observed value (for<br>SHV>median) | P-Value |
| --- | --- | --- | --- | --- |
| sum(D - B) | -15.6 % | 0.0015 | 16.1 % | 0.0006 |
| sum((D - C) - (B - A)) | -28.6 % | 0 | 29.1 % | 0 |
| sum(D - C) | -13.1 % | 0.0025 | 13.15 % | 0.0021 |
| sum(B - A) | 15.5 % | 0.0011 | 16.0 % | 0.0008 |

For the precision analysis we used the coefficient of variation (equation 1) as a metric to show the precision level by calculating the ratio of standard deviation (equation 2) over the mean of N joint synapses. Here N is 2 but can take higher values as up to 7 have been detected in previous studies. Since this is a sample from the unknown population of joint synapses, we used the corrected standard deviation formula with 1/(N-1) factor.

The coefficient of determination, denoted *R*^2^, was used in Fig. 4 panel A-E to show the proportion of the variation in the dependent variable (CV) that is predictable from the independent variable (spine head volumes).

One factor Kruskal-Wallis (KW) test was in ***Fig. 4*** F to check for a significant difference between the 4 dentate gyrus SDSA datasets and the CA1 dataset.

Logarithmic scale was used on skewed distributions (Fig. 2 A-F. Fig. 4, A-E. Fig. 5 B and C. Fig. 6 A-D.)

Boxplots in ***Fig. 4*** F are made as follows: The lower and upper hinges correspond to the first and third quartiles (the 25th and 75th percentiles). The upper whisker extends from the hinge to the largest value no further than 1.5 * IQR from the hinge (where IQR is the interquartile range, or distance between the first and third quartiles).

Bootstrapping was done by combining algorithms 1 and 2 to calculate the standard errors as explained below in the sections *Standard error of Median* and *Clustering Algorithm*.

The standard errors of the entropy, efficiency constant, maximum entropy for uniform distribution, and KL divergence (Table 2 column 2-5) are all calculated similarly using the bootstrapping technique explained in algorithm 1. (See (***Efron et al., 2021***) for further information regarding bootstrapping for the calculation of standard error.)

### Standard error of Median

The standard error of median for the precision levels of each of the 5 datasets’ SDSA pairs is calculated with Algorithm 1 as follows. The idea is to generate 1000 bootstrap samples of length n, each sampled from the n SDSA pairs with replacement, to estimate the standard error of median for the n SDSA pairs (Table 1, column 3). The standard error of median of spine head volumes follows the same procedure using Algorithm 1.

#### Algorithm 1 Bootstrap Algorithm for Estimating the Standard Error of Median

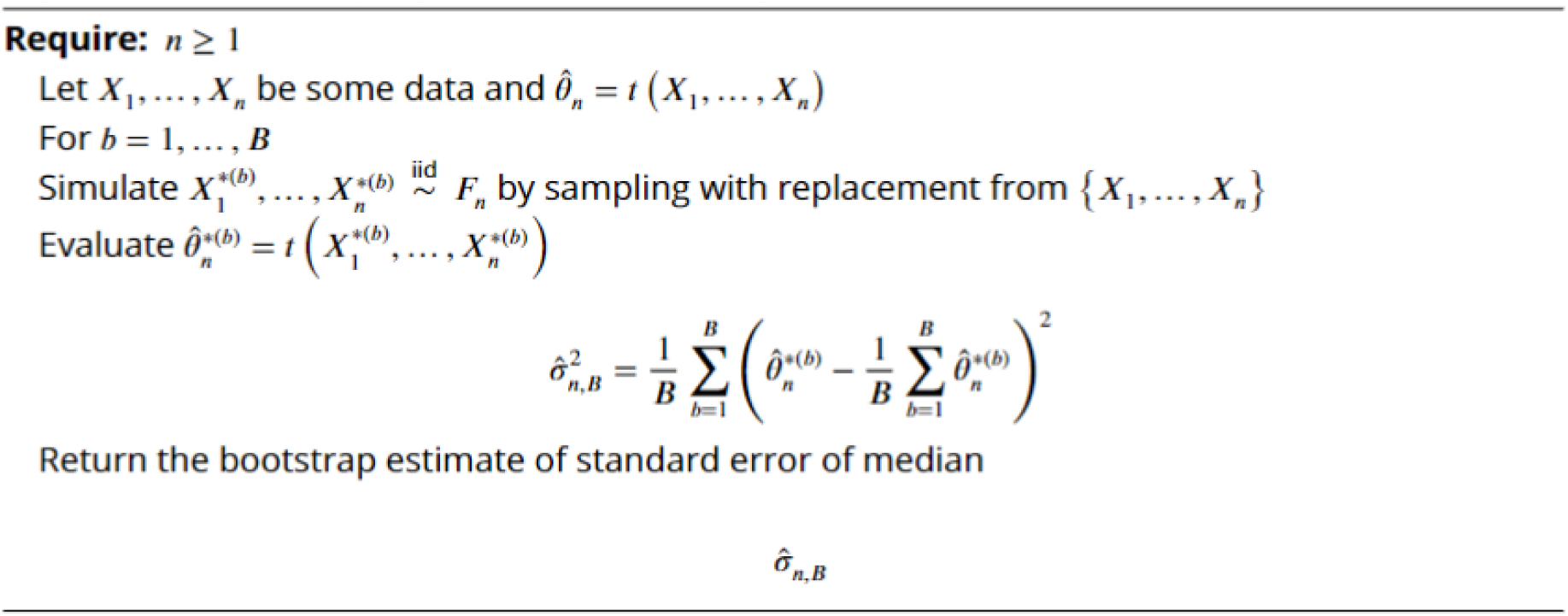

### Clustering Algorithm

To construct the clusters, spine head volumes are sorted from smallest to the largest. The first value (smallest value) is selected and the CV of that value and the remaining head volumes are calculated in a pairwise manner. The head volumes for which the calculated CV is below the threshold (the median value of the SDSA pairs CV) are assigned to the first category and deleted from the pool of *N* spine head volumes. This procedure is repeated until the CV exceeds the median SDSA pairs CV and a new category is formed. New categories are formed until all the remaining spine head volumes are assigned to a category and the original vector of spine head volumes is empty (see Algorithm 2 for details). It is guaranteed that the coefficient of variation between each random pair within each category is less than the threshold value measured from the reconstructed tissue SDSA pairs. All spine head volumes are rounded to two significant digits for the display.

#### Algorithm 2 Clustering Algorithm

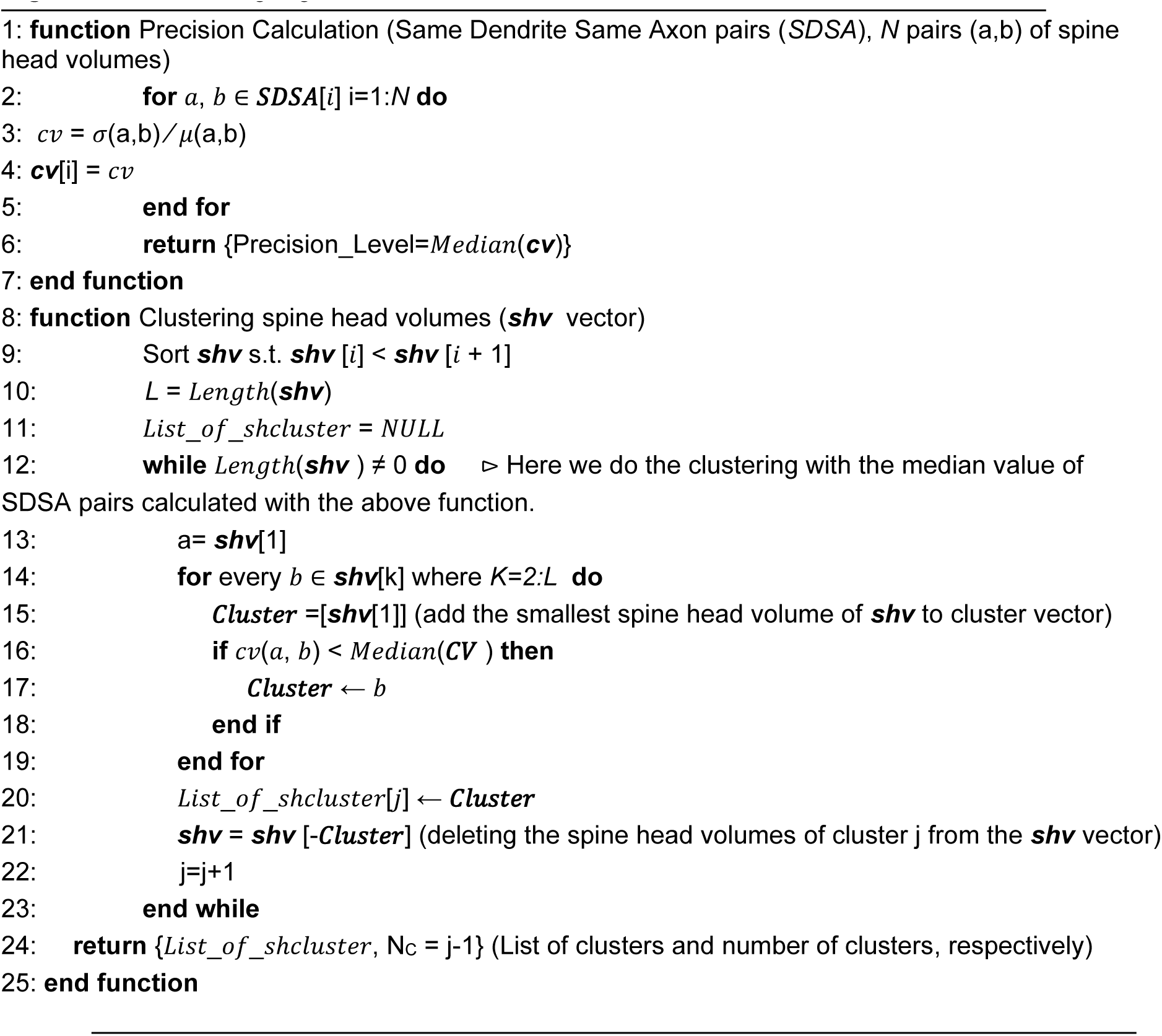

For each panel in Fig. 6, the Y axis shows the percentage of spine head volume counts in the respective bin. The area under each plot is normalized to *1* for a fair comparison. The X axis shows the spine head volumes in *µm*^3^ on the log scale. The width of each bin is exactly the median value of the set of CV values for each condition calculated in Fig. 5 (Example: for bin-1 panel-1 [x1,x2], CV(x1,x2)=0.65, where x1 is the smallest spine head volume in dentate gyrus 30 min control dataset and x2 is a larger hypothetical head volume that has a CV of 0.65 with x1). The height of bin 1 is the number of spine head volumes in that range normalized to the total number of spine head volumes in that dataset (Example: for Fig. 6A for dentate gyrus 30 min control it is 236). (A) dentate gyrus 30 min control. (B) dentate gyrus 2hr LTP. (C) dentate gyrus 30 min LTP. (D) dentate gyrus 2hr LTP.

### Information and Entropy in Synaptic Plasticity

“The fundamental problem of communication is that of reproducing at one point either exactly or approximately a message selected at another point. Frequently the messages have meaning; that is they refer to or are correlated according to some system with certain physical or conceptual entities. These semantic aspects of communication are irrelevant to the engineering problem. The significant aspect is that the actual message is one selected from a set of possible messages. The system must be designed to operate for each possible selection, not just the one which will actually be chosen since this is unknown at the time of design.” (<u>Shannon</u>***, 1948)***

Shannon’s information theory is the rigorous basis for a new method to quantify empirically SISC; that is, the number of bits of Shannon entropy stored per synapse. For this new method, only the precision as measured by the coefficient of variation (CV) of SDSA pairs, illustrated in Fig. 4, was borrowed from Bartol et al. (2015). The new method performs non-overlapping cluster analysis (Fig. 5 and 6) using Algorithm 2, to obtain the number of distinguishable categories of spine head volumes from the CV measured across the reconstructed dendrites.

The Shannon information per synapse is calculated by using the frequency of spine head volumes in the distinguishable categories where each category is a different message for the calculation of Shannon information. The maximum number of bits is calculated as the *log2(N_C_)* where *N_C_* is the number of categories which set an upper bound for the SISC.

When calculating the amount of entropy per synapse, the random variable is the synapse size and the number of distinguishable synaptic states is the realization of a random variable for the occurrence of each state. The probability of the occurrences of each state is calculated by the fraction of the number of spine head volumes in each of the clusters over the total number of spine head volumes in the reconstructed volume.

The information coding efficiency at synapses is measured by Kullback-Leibler (KL) divergence to quantify the difference between two probability distributions, one from the categories of spine head volumes and the other from a corresponding uniform distribution with the same number of categories.

### Synaptic Information Storage Capacity

Spine morphology has substantial variation across the population and lifetime of synapses. Hebbian plasticity puts forth a causal relationship and transformation of information from the presynaptic site to the postsynaptic site by the adjustment of efficacy of synaptic transmission, or “synaptic weight.” The pattern of synaptic weights in the ensemble of neural circuits allows us to define both information and the recipient of the message in the context of synaptic plasticity. The recipient of the message is the neural ensemble or the pattern of synaptic weights that store the message and read the message during the recall process, which is the reactivation of the synaptic weights in the memory trace. The amount of information is quantified by the distinguishability of synaptic weights which comprise the memory trace. Here “distinguishability” implies that the precision of synaptic weights play a significant role.

The synapse is the unit of information storage in an ensemble of neurons, and if the precision level of synaptic weights is low then the amount of information that can be stored per synapse and in the ensemble of the neurons will also be low. Because the spine head volume is highly correlated with synapse size, the precision of spine head volumes can be used to measure the distinguishability of the synaptic weights. High precision yields a greater number of distinguishable categories (i.e., states or clusters) for spine head volumes and hence higher information storage capacity.

## Acknowledgments

We would like to acknowledge Adel Aghajan and Wenxin Zhou for helpful discussion regarding information theory and bootstrapping analysis and Patrick Parker for editorial support. This research was supported by NSF 170756; NSF 2014862; NIH P41GM103712; NIH R01 - NIH MH095980-07; NIH MH115556; NIH MH129066.

## Author contributions

M.S., T.M.B., W.C.A., K.M. H, and T.J.S. designed research; M.S., T.M.B., W.C.A., K.M.H., and T.J.S., analyzed data; M.S., designed and implemented all simulation algorithms and generated results for the manuscript and applied the information theory to the analyses with contributions from T.M.B., W.C.A., K.M.H. and T.J.S.; M.S., T.M.B., J.B.B., D.D.H., D.C.H., P.V.G., M.K., J.M.M., P.H.P., W.C.A., K.M.H, and T.J.S. performed research; J.B.B., M.K., J.M.M., W.C.A., and K.M.H. designed and performed the electrophysiology experiments, tissue processing, and imaging; M.S., C.B., J.B.B., D.D.H., D.C.H., P.V.G., M.K., P.H.P., and K.M.H. performed, curated reconstructions; M.K. and D.D.H. made martials for Fig. 1; M.S. and T.J.S. wrote the paper with contributions from T.M.B., W.C.A., K.M.H. This research is part of multi-institutional collaboration project NeuroNex 1 & 2 led by K.M.H.

---

**Supplementary Fig 1.**
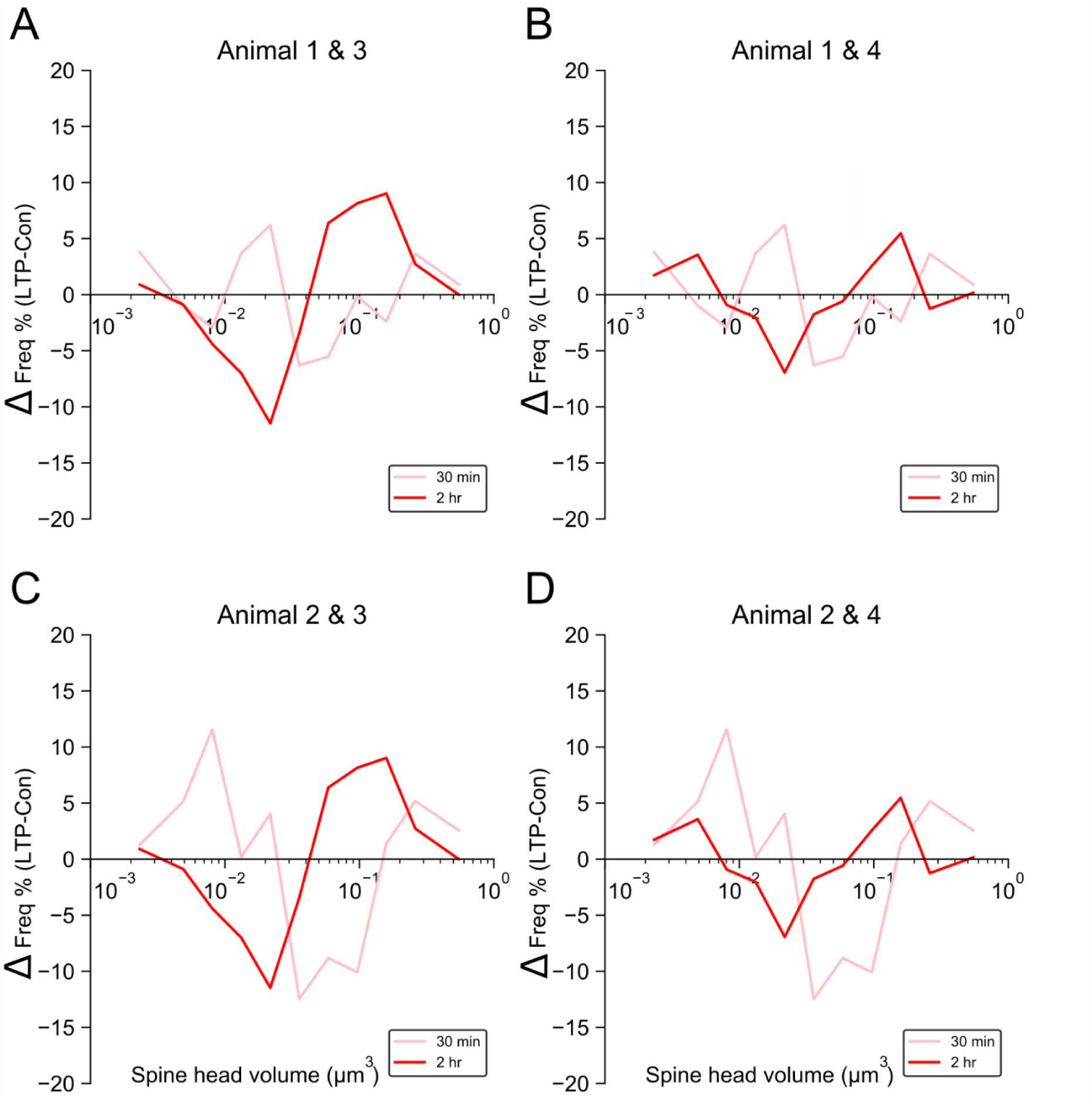
Change relative to control hemispheres in the distribution of spine head volumes at 30 min and 2 hr after the induction of LTP per animal. Difference between the frequency of spine head volumes in 30 min LTP and 2 hr LTP conditions. (A) Animal 1 & 3 (B) Animal 1 & 4 (C) Animal 2 & 3 (D) Animal 2 & 4.

**Supplementary Fig 2.**
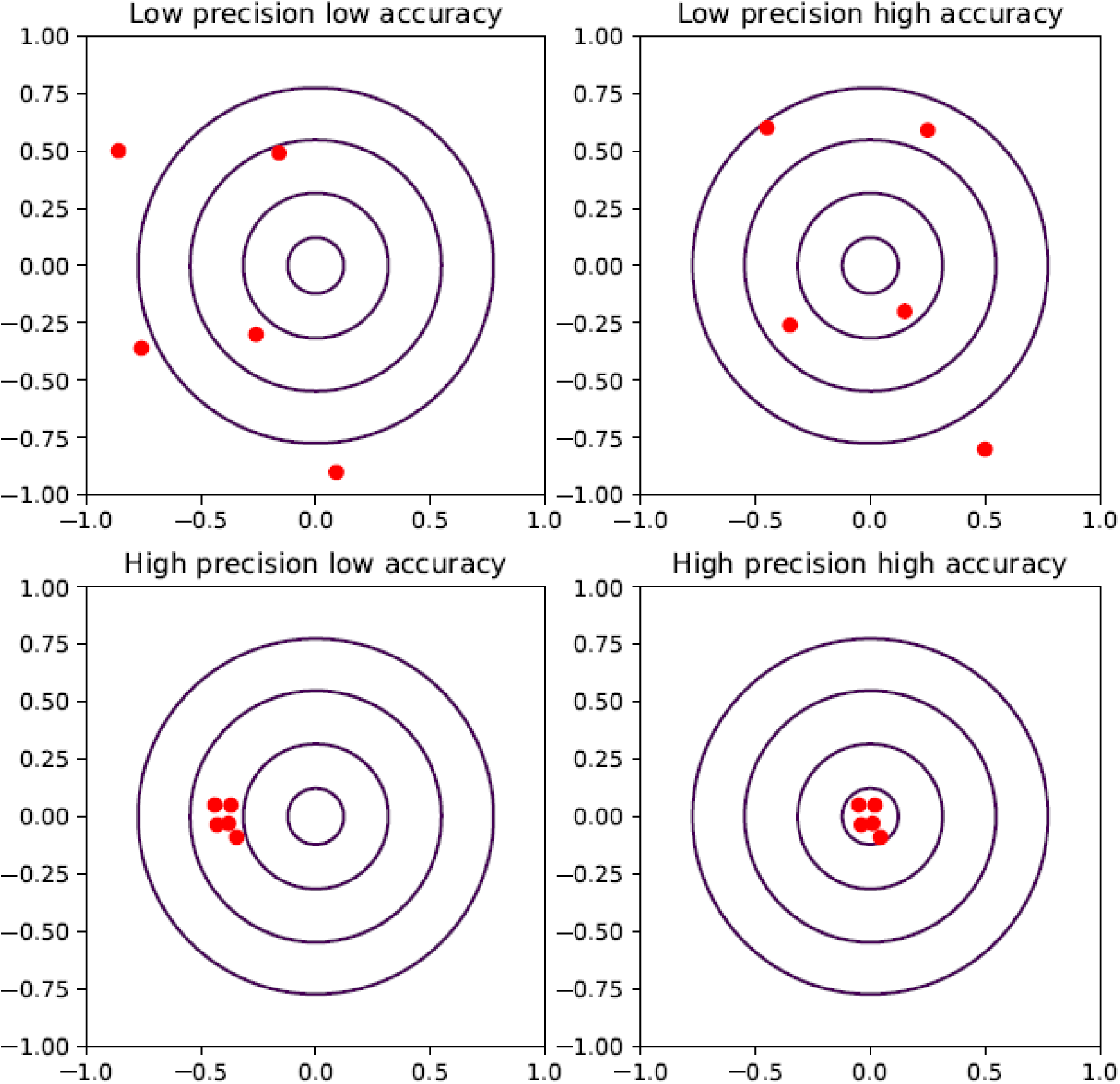
Dart precision versus accuracy. Precision concerns the degree of reproducibility of a process. When a process or system is repeated with the same input the amount of variation in the output shows the precision level of the process. For accuracy there is a reference frame with which the average value of measurements is compared. The graphs illustrate a low precision and low accuracy outcome (top left), low precision and high accuracy (top right; the average of the positions is almost on the bull’s eye), high precision and low accuracy (bottom left), and high precision and high accuracy (bottom right).

**Supplementary Fig 3.**
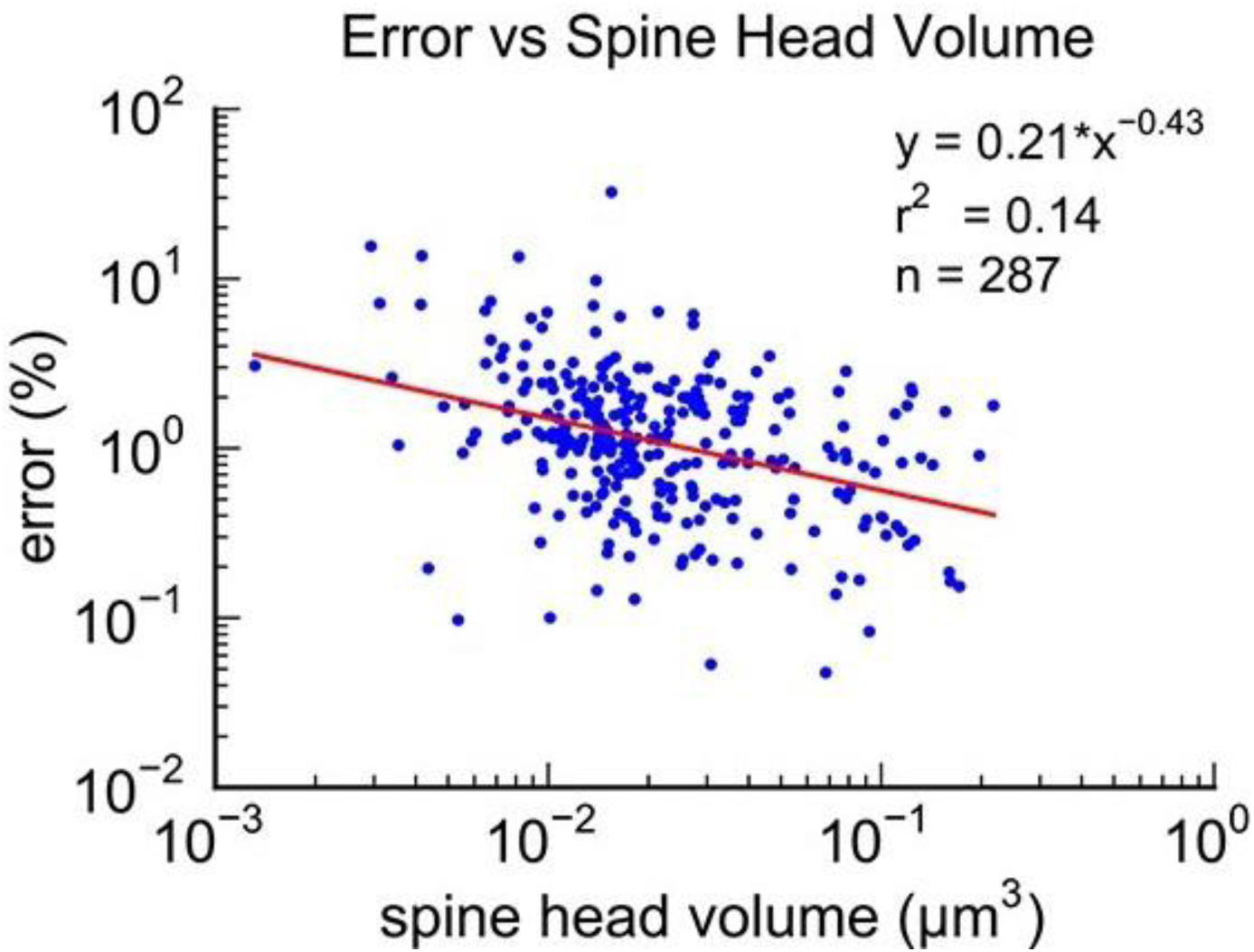
The 3D reconstructions of the spine head volumes of the 5 datasets were performed with the same protocol as that used for the CA1 dataset in ***Bartol et al., eLife, (2015)***. Four individuals made hand tracings from the 2 dimensional electron micrographs and then after alignment the automatic 3D reconstruction was made. The average measurement error is about 0.01 as shown in the above figure.

**Supplementary Fig 4.**
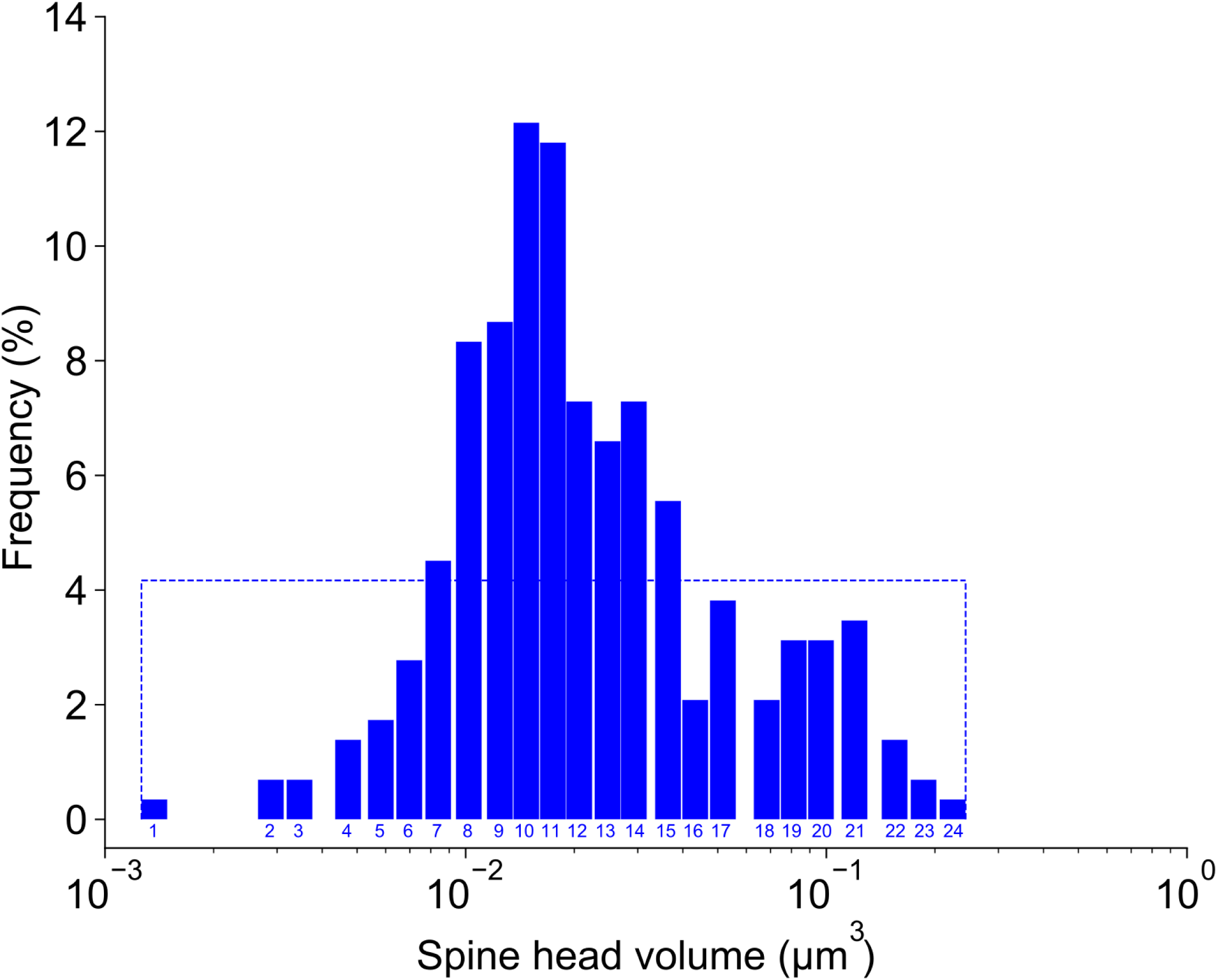
CA1 24 distinguishable clusters. The Y axis indicates the frequency of spine head volumes within each cluster and the X axis indicates the spine head volumes values in the log scale. The dash rectangular box around the histogram is the frequency of spine head volumes if the 288 spine head volumes were distributed uniformly among the 24 clusters.

**Supplementary Fig 5.**
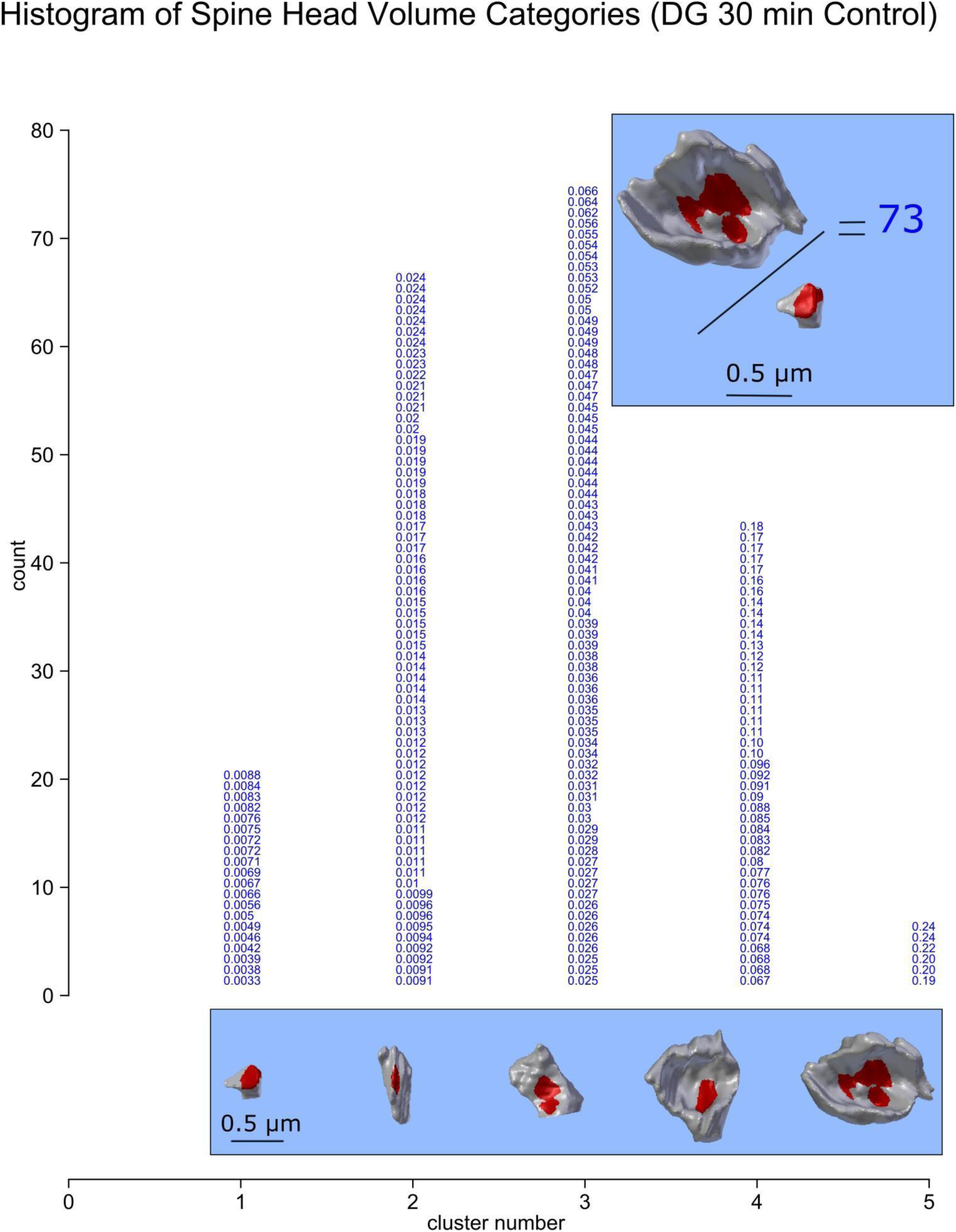
The clustering of 209 spine head volumes of two rats in control conditions (30 min data). To analyze synapses in the whole reconstructed cube, the 209 spine head volumes are clustered into 5 distinguishable categories based on the median value CV calculated from 10 SDSA pairs detected in the reconstructed cube. Median value of 10 CVs calculated from analysis illustrated in inset in panel B with the value of 0.65. The Y axis shows the number of spine head volumes within each category. The 3D object below each category (vertical column) is the actual 3D reconstructed spine head volume of the largest head volume in the category. The X axis shows the distinguishable categories number.

**Supplementary Fig 6.**
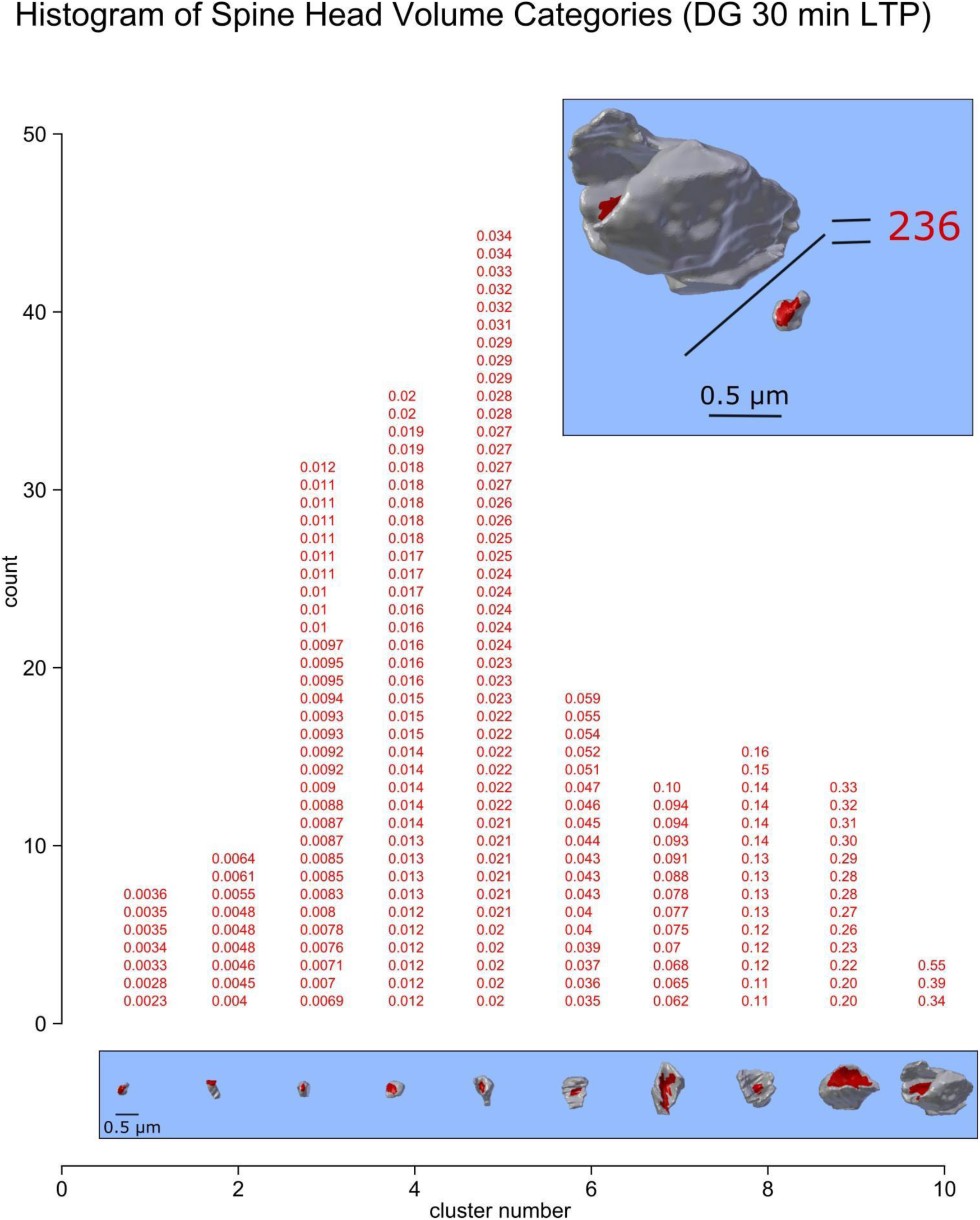
The clustering of 188 spine head volumes of two rats in LTP conditions (30 min data).

**Supplementary Fig 7.**
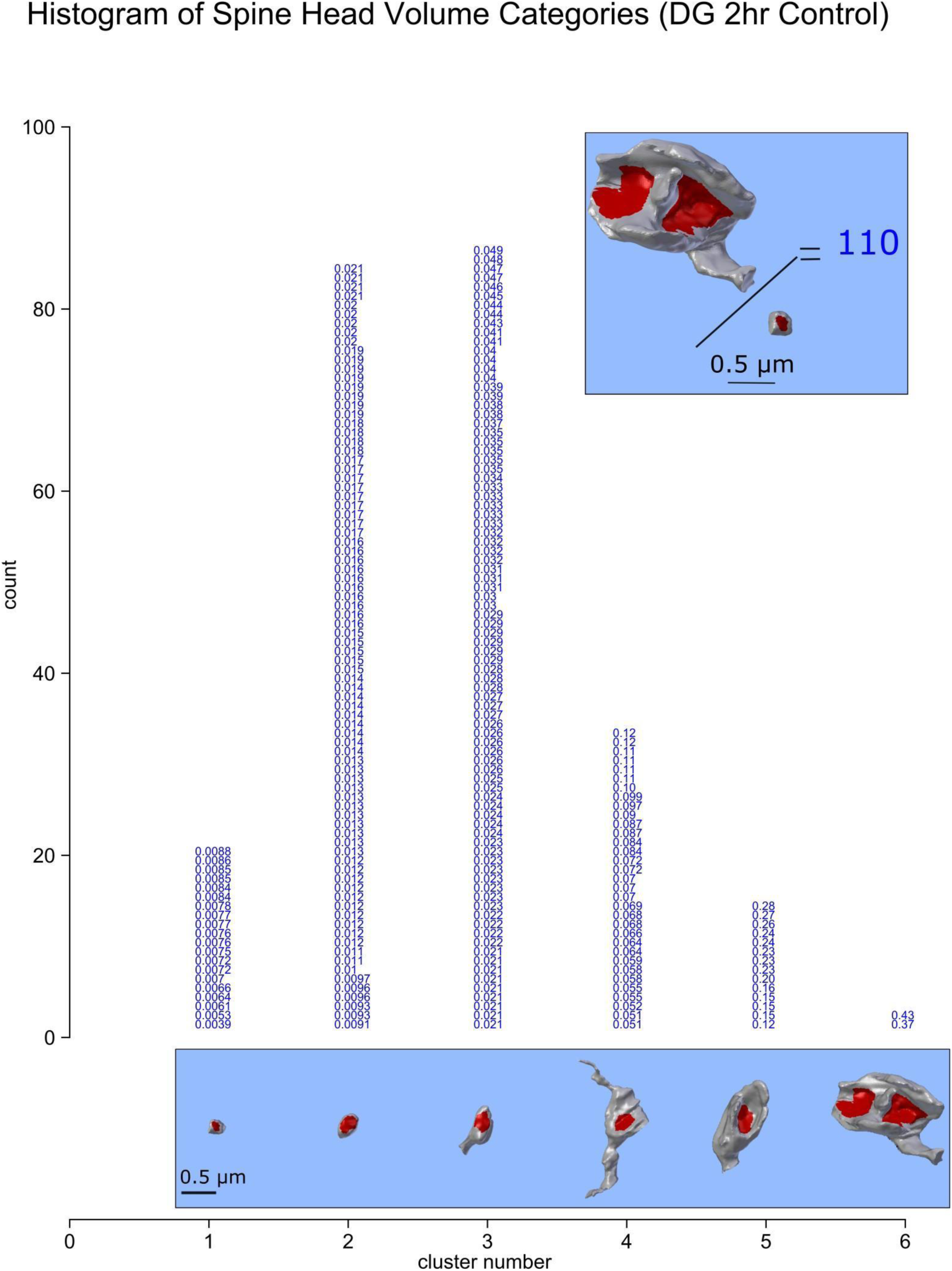
The clustering of 239 spine head volumes of two rats in control conditions (2 hr data).

**Supplementary Fig 8.**
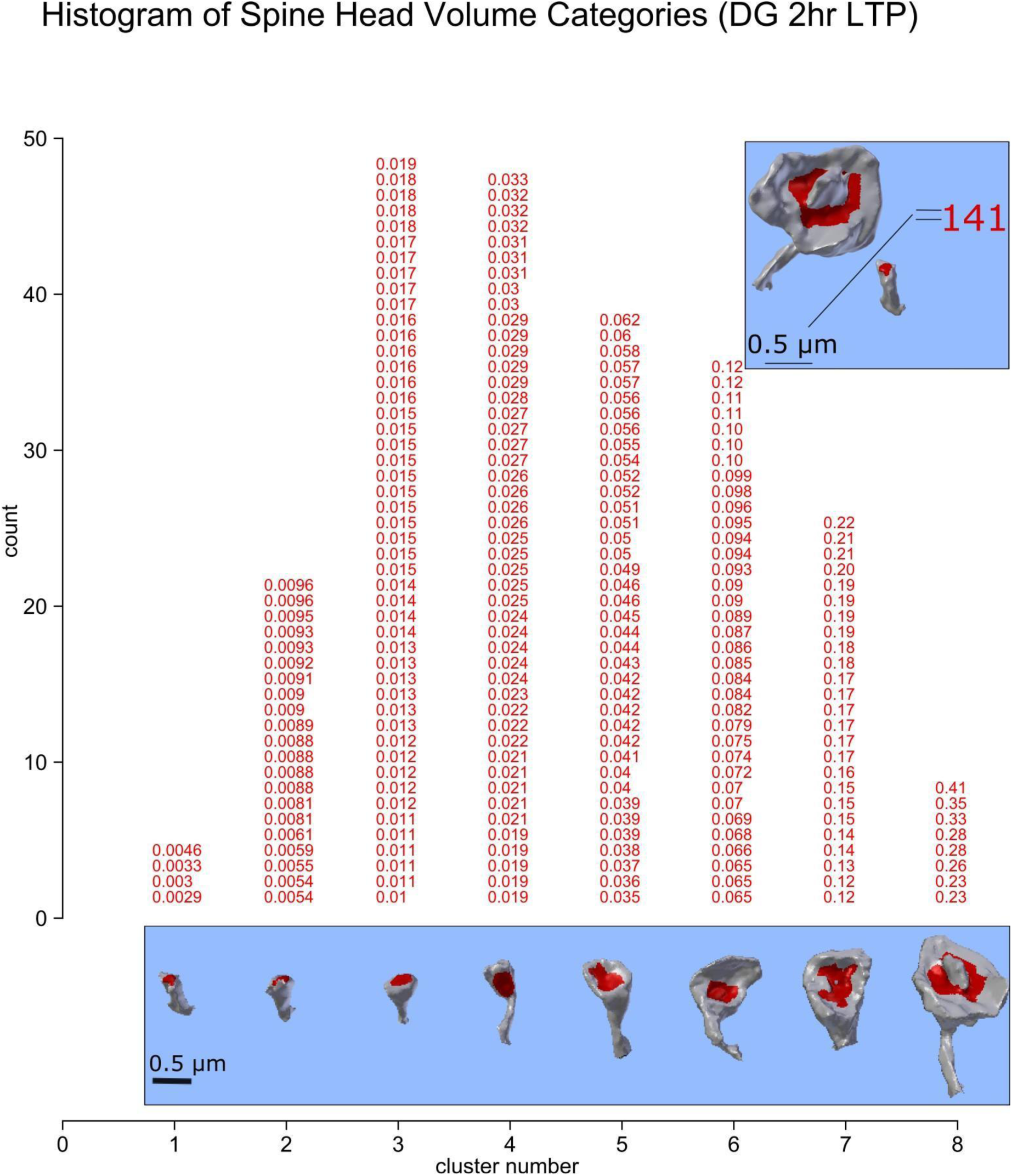
The clustering of 226 spine head volumes of two rats in LTP conditions (2 hr data).

